# BMP signaling during gastrulation pre-patterns the dorsal spinal cord

**DOI:** 10.64898/2026.05.18.726076

**Authors:** Hannah Greenfeld, Daniel E. Wagner

**Affiliations:** Department of Obstetrics, Gynecology and Reproductive Science, Center for Reproductive Sciences, University of California San Francisco; San Francisco, USA; Eli and Edythe Broad Center for Regeneration Medicine and Stem Cell Research of Obstetrics, Gynecology and Reproductive Science, Center for Reproductive Sciences, University of California San Francisco; San Francisco, USA

## Abstract

The classic model of dorsal spinal cord patterning proposes that roofplate-derived BMP patterns dorsal interneuron subtypes in a concentration-dependent manner. However, genetic perturbations of BMP pathway components produce variable effects, challenging this model. Here we implemented single-cell profiling, fate mapping, and mosaic perturbations to determine when BMP signaling patterns dorsal neural fates *in vivo*. Contrary to the classic model, we demonstrate that dorsal fates are patterned by BMP signaling during gastrulation. Following neural tube formation, BMP signaling continues but plays limited roles in domain specification and maturation. Fate mapping revealed that dorsal progenitors originate from the ventral gastrula, adopting BMP-dependent transcriptional states that prime dorsal neural fate. We propose that dorsal neural fates are initially patterned by gastrulation-stage sources of BMP, prior to roofplate induction.

## Introduction

The vertebrate neural tube has served as a paradigm for understanding how morphogen gradients pattern tissues and specify cell types throughout development. Within the neural tube, the emerging central nervous system is organized into functionally distinct neuronal subtypes whose positions along the dorsoventral axis are mirrored by opposing morphogen gradients^1^. The classic model for this process holds that graded Sonic hedgehog (Shh) from the ventral floorplate and Bone Morphogenetic Protein (BMP) from the dorsal roof plate act in concentration-dependent fashion to instruct progenitor fate^2,3^. Specifically in dorsal regions, BMP signaling from the roof plate is proposed to pattern dI1–dI3 subtypes in a concentration-dependent manner, with higher levels specifying more dorsal identities^4–6^. However, genetic perturbations of BMP pathway components produce surprisingly selective and inconsistent phenotypes that are difficult to reconcile with this simple morphogen model^4,7–10^. Critically, most studies have also focused on BMP activity following neural tube closure, leaving unexplored whether earlier BMP signaling during gastrulation might contribute.

Here, we use the zebrafish embryo to resolve the *in vivo* time windows in which BMP signaling patterns dorsal neural fates. Using single-cell transcriptomics, time-resolved BMP inhibition, mosaic perturbations, and photoconversion-based lineage tracing, we reveal that dorsal spinal cord patterning occurs through a multi-phase mechanism in which gastrulation-stage BMP signaling pre-patterns progenitor competence, while later BMP activity within the neural tube refines a single subtype’s specification and neuronal maturation.

## Results

### Single-cell transcriptomics reveal molecularly distinct but spatially intermixed dorsal interneuron subtypes

To define the cellular composition of the zebrafish dorsal spinal cord and the role of BMP signaling in neural patterning, we generated a single-cell atlas of interneuron cell types within the dorsal neural tube (dNT). We performed single-cell RNA sequencing (scRNA-seq) on dissected trunks from 48-hours post fertilization (hpf) embryos treated with either DMSO or 50μM DMH1, a small molecule BMP type 1 receptor inhibitor^11^ (Fig. 1a). The resulting scRNA-seq map revealed six molecularly distinct dorsal interneuron (dI) subtype clusters (dI1–6) within the dorsal spinal cord (Fig. 1b), with subtype identities confirmed by marker-gene scoring against established transcription factor combinations (Fig. 1c; Supplementary Fig.1a-e)^1,6,12^.

**FIGURE 1.**
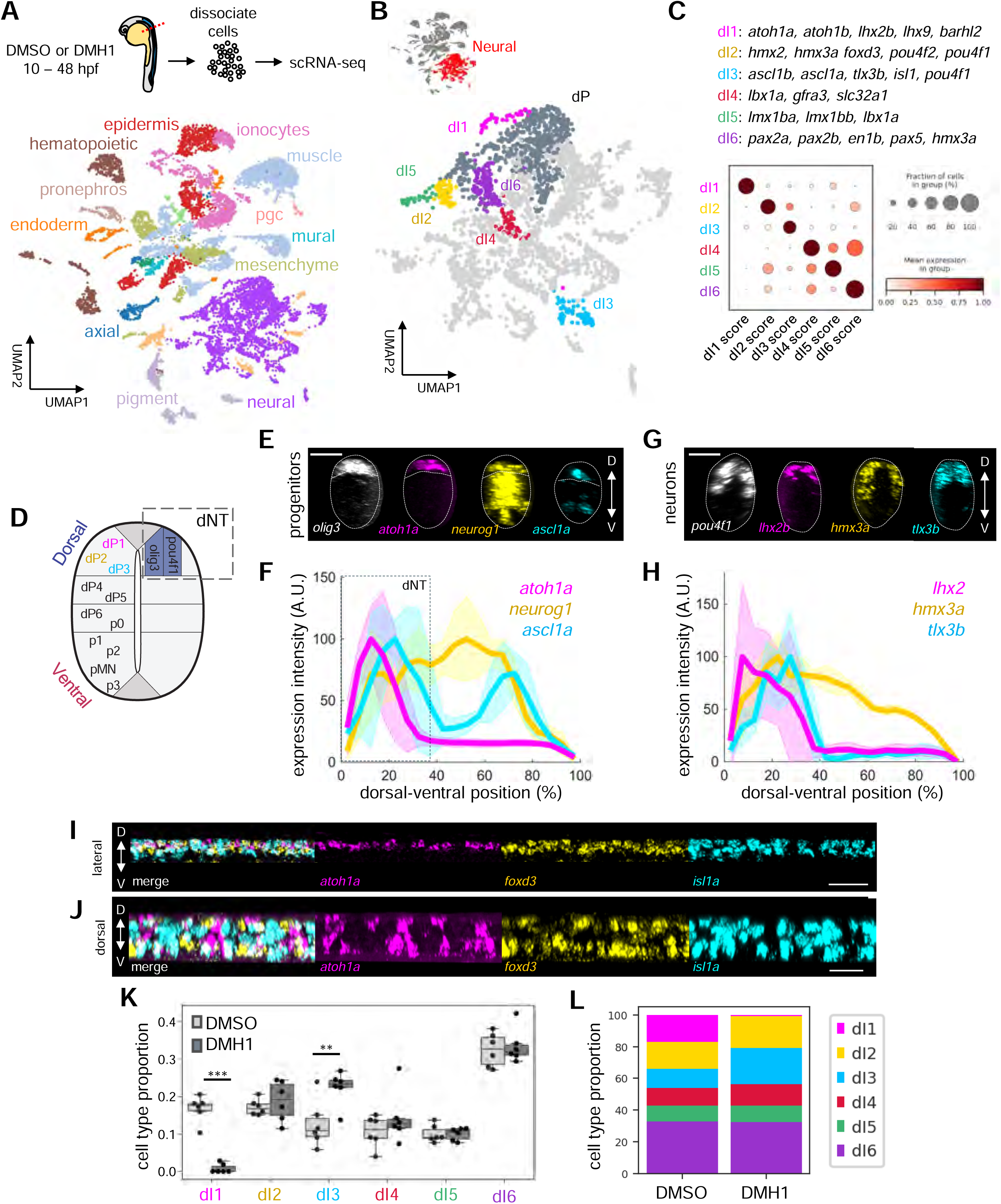
Zebrafish dorsal interneurons are molecularly distinct but spatially intermingled. **a** Top, experimental schematic for single-cell RNA sequencing. Trunks from 48 hpf embryos treated with either DMSO or the BMP type I receptor inhibitor DMH1 (10–48 hpf) were dissected and dissociated for scRNA-seq. Bottom, Uniform Manifold Approximation and Projection (UMAP) plot of scRNA-seq (nCells = 11,942) from 48 hpf zebrafish trunks, colored by cell type annotation. **b** Subset of UMAP plot of annotated neural cell types. Dorsal interneuron subtypes (dI1–6) are highlighted. **c** Dotplot of mean gene expression score for each dI cluster based on canonical transcription factor markers. Marker genes used to generate the scores are listed above. Each row represents a cell cluster; each column represents a marker gene score. **d** Schematic of the dorsal neural tube (dNT) marked by expression of *olig3* in dorsal progenitors and *pou4f1* in dorsal neurons dI1-3. **e** Representative images for orthogonal views of HCR FISH at 48 hpf for all dorsal progenitors (*olig3*), dP1 (*atoh1a*), dP2 (*neurog1*), and dP3 (*ascl1a*). Dorsal is up. Scale bar, 25 μm. **f** Normalized fluorescence intensities of markers for dorsal progenitors dP1 (*atoh1a*), dP2 (*neurog1*), and dP3 (ascl1a) across the DV axis of the neural tube at 48 hpf (dorsal = 0%, ventral = 100%). Box marks region of the dNT. n = 5, each). **g** Representative images for orthogonal views of HCR FISH at 48 hpf for all dorsal neurons (*pou4f1*), dIN1 (*lhx2b*), dIN2 (*hmx3a*), and dIN3 (*tlx3b*). Dorsal is up. Scale bar, 25 μm. **h** Normalized fluorescence intensities of markers for dorsal interneurons dIN1 (*lhx2b*), dIN2 (*hmx3a*), and dIN3 (tlx3b) across the DV axis of the neural tube at 48 hpf (dorsal = 0%, ventral = 100%). Box marks region of the dNT n = 5, each). **i-j** Representative images for lateral views (i) and dorsal views (j) of HCR FISH at 48 hpf for all dorsal interneurons dI1 (*atoh1a*), dI2 (*foxd3*), and dI3 (*isl1a*). Scale bar, 30 μm. **k** Analysis of differential cell type abundance (see Methods) of dorsal interneurons between DMSO- and DMH1- treated embryos. ***FDR <0.01, **FDR <0.05. **l** Stacked bar plot of dorsal interneuron subtype composition in DMSO- and DMH1- treated embryos.

To determine how these dorsal cell types (marked by *olig3*+ progenitors and *pou4f1*+ neurons^13–15^) are organized within the neural tube, we performed HCR fluorescent *in situ* hybridization (FISH)^16^ using additional specific markers of the three most dorsal dI subtypes (Fig 1d). Both progenitor and differentiated dorsal interneuron markers were predominantly expressed in the dorsal region of the neural tube (Fig. 1e-h). These data also revealed that distinct progenitor subtypes were spatially intermixed at similar dorsoventral levels, rather than occupying discrete domains^1,17^ (Fig. 1i, j, Supplementary Fig.1f). Such patterns are difficult to reconcile with the classic model, which predicts domain-level specification as a function of distance from the roofplate. This finding thus prompted us to reconsider the mechanism of dorsal progenitor specification.

### BMP signaling within the neural tube is dispensable for specification of most dorsal progenitors

We next explored how loss of BMP signaling affected dI subtype composition. Treatment of embryos with 50μM DMH1 from 10-48 hpf abolished phosphorylation of the BMP transcriptional effector Smad1/5/9 (pSmad1/5/9) across the dorsoventral axis of the neural tube (Supplementary Fig. 2a, b). Consistent with a prior scRNA-seq analysis of BMP pathway disruption^18^, cells from both conditions were present across all clusters, indicating BMP inhibition did not broadly disrupt cell identity (Supplementary Fig. 2c). Across all embryonic tissues, DMH1-dependent cell type composition changes mirrored the expected dorsalized phenotype of BMP loss-of-function^19^. Known BMP-dependent tissues, e.g. fin epidermis, were significantly reduced in DMH1-treated embryos, consistent with effective pathway inhibition (Supplementary Fig. 2d, Supplementary Table 1). Within neural tissues, differential abundance analyses revealed a striking and selective effect inconsistent with the classic model: only dI1 neurons were significantly reduced in DMH1-treated embryos (Fig. 1k). By contrast, proportions of dI2 neurons were unchanged, dI3 neurons increased in abundance, and the overall proportion of dI4-6 neurons remained unaffected. Analysis of subtype composition indicated that dI3 expansion likely occurred at the expense of the dI1 population (Fig. 1l).

That dI2 and dI3 neuron clusters appeared unchanged and increased in abundance, respectively, following the loss of BMP signaling represented a striking departure from predictions of the classic model. To validate these results, and to determine whether compositional changes were accompanied by spatial reorganization, we performed HCR FISH for markers of dI1–3 neurons (Supplementary Fig. 3a). Consistent with our scRNA-seq results: (1) expression of multiple dP1 progenitor (*atoh1a*, *atoh1b*) and dI1 neuronal (*lhx2*) markers were lost following BMP inhibition (Fig. S3b-d, g); (2) marker expression for dI2 progenitors (*neurog1*) and early-stage neurons (*lhx1*, *hmx2*) remained unchanged (Supplementary Fig. 3b, c, e, g); and (3) expression of dI3 markers (*ascl1*, *tlx3b, hs3st1l2*) increased – and shifted dorsally (Supplementary Fig. 3b, c, f, g). In addition, we also noted diverging trends for dI2 markers with different temporal dynamics: *lhx1a*, expressed early during neuronal differentiation, and *foxd3*, activated only upon maturation^20–22^. Across both HCR spatial profiling and scRNA-seq, DMH1 treatment led to a specific elimination of dorsal *foxd3* expressing cells (with ventral *foxd3* marking V1 neurons unaffected) while leaving *lhx1a* expression and overall dI2 neuron cluster size unaffected (Supplementary Fig. 3g, Supplementary Fig. 4a-e). Thus, dI2 neurons are specified independently of BMP signaling but require it to activate the full transcriptional program of mature dI2 identity.

Together, these findings indicate that BMP signaling within the neural tube is specifically required for dI1 specification as well as for the maturation, positioning, and/or restriction of additional dorsal fates. However, roofplate-associated BMP signaling is apparently not required for the specification of dI2 or dI3 progenitor subtypes, as predicted by the classic morphogen model.

### BMP signaling during gastrulation is required for dorsal neuron specification

Given that BMP signaling from 10-48 hpf was not required to specify most dorsal cell fates, we next asked whether BMP signaling at an earlier developmental stage might instead pre-pattern the dorsal neural tube. During gastrulation, the embryo is exposed to graded BMP signaling^23^, raising the possibility that this earlier signal establishes dorsal progenitor identity. To avoid the embryonic lethality caused by global inhibition of BMP signaling during gastrulation, we leveraged mosaic embryos to test whether rare cell clones subject to autonomous BMP signaling inhibition would be competent to adopt later dorsal interneuron fates. We thus generated mosaic BMP-inhibited embryos by co-injecting one blastomere of 8-cell stage embryos with transposase mRNA and a plasmid containing a Tol2-flanked heat-shock inducible dominant-negative BMP type 1 receptor (dnBMPR1)^24^ together with a constitutively expressed H2B-NeonGreen lineage tracer. The mosaic and inducible nature of the dnBMPR1 expression enables embryos with these perturbed clones to complete gastrulation and develop to 48 hpf with normal morphology (Fig. 2a).

**FIGURE 2.**
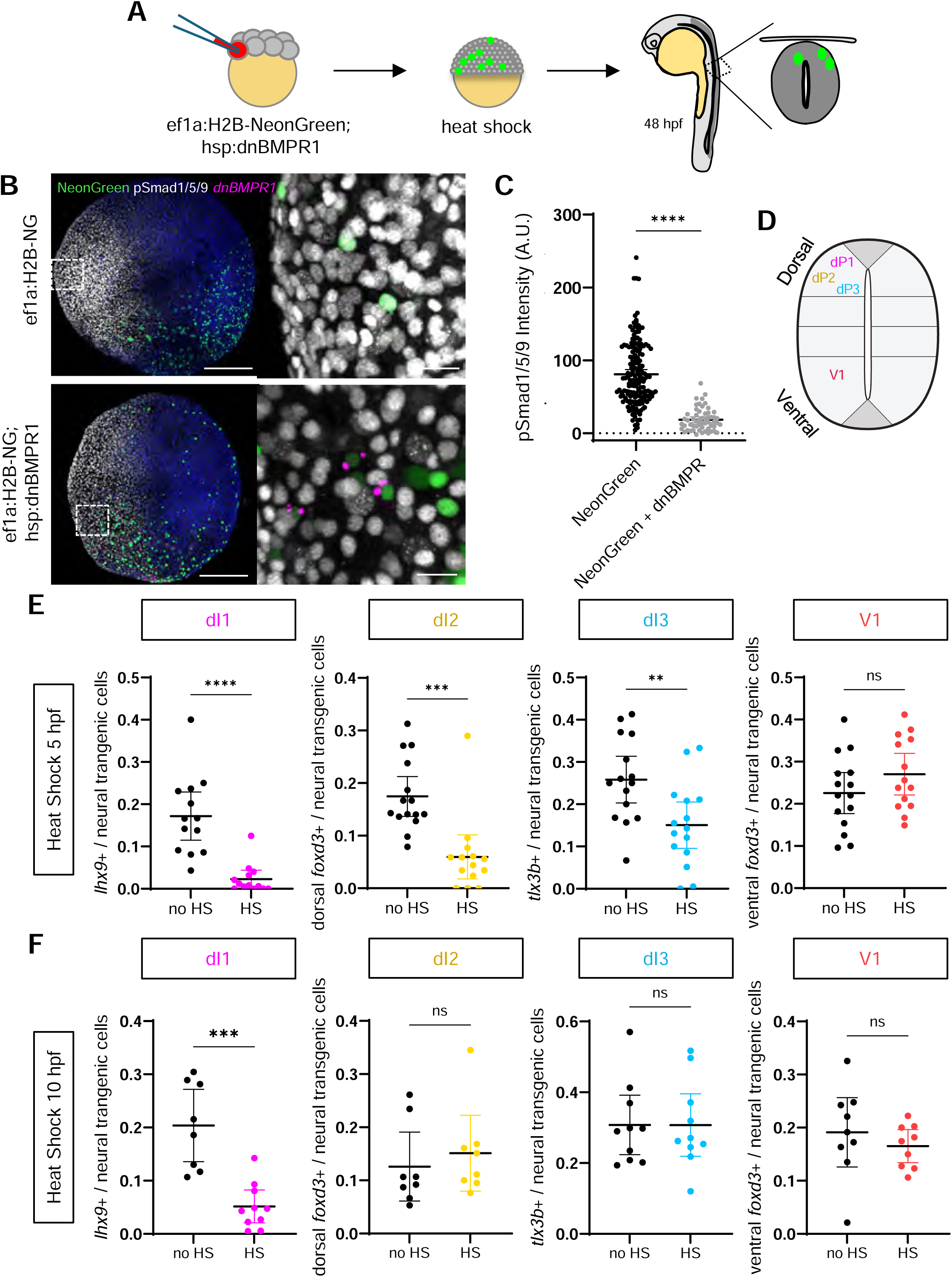
BMP signaling during gastrulation is required for dorsal neuron specification. **a** Experimental schematic of cell-autonomous inhibition of BMP signaling activity by mosaic analysis of Tg(*hsp*:dnBMPR; *eef1a1l1*:h2b-NeonGreen) or Tg(*eef1a1l1*:h2b-NeonGreen). Tol2 mRNA and plasmid DNA were co-injected into one blastomere of an 8-cell stage embryo. Embryos are heat-shocked at 5 or 10 hpf to induce expression of dnBMPR and transgenic cells are evaluated at 48 hpf for specification into specific dorsal neurons. **b** Representative images of pSmad1/5/9 immunostaining and dnBMPR1 FISH of wild-type embryos injected with either *eef1a1l1*:NeonGreen alone or *eef1a1l1*:NeonGreen; hsp:dnBMPR1 and fixed 2 hours after heat-shock at 5 hpf. Scale bars, 200 μm and 20 μm (inset). **c** Quantification of nuclear pSmad1/5/9 fluorescent intensities of cells expressing either *eef1a1l1*:NeonGreen alone or *eef1a1l1*:NeonGreen; hsp:dnBMPR1 fixed 2 hours after heat-shock. (n=5). **d** Schematic of the dorsal cell types: dI1, 2, 3, and the V1 neuron within neural tube. **e-f** Proportion of transgenic cells co-expressing *dnBMPR1* and markers for dIN1 (*lhx9*), dIN2 (dorsal *foxd3*), dIN3 (*tlx3b*), and V1 (ventral *foxd3*) from all transgenic cells in the neural tube after heat-shock at 5 hpf (D) or 10 hpf (E). ****P<0.0001, ***P<0.001, **P<0.01.

To confirm effective inhibition of BMP signaling during gastrulation, we heat-shocked mosaic embryos at 5 hpf and assessed pathway activity by staining for pSmad1/5/9 two hours later (Fig. 2b). Cells expressing dnBMPR1 and NeonGreen showed a significant reduction in pSmad1/5/9 levels compared to cells expressing NeonGreen alone, confirming efficient cell-autonomous inhibition of BMP signaling in marked cells (Fig. 2c). We next asked whether cells unable to transduce BMP signaling during gastrulation could later adopt dorsal interneuron identities (Fig. 2d). Quantification of transgenic cells co-expressing dnBMPR1 and subtype-specific neuronal markers at 48 hpf revealed a significant reduction in the proportion of labeled cells differentiating into *lhx9*+ dI1, *foxd3*+ dI2, and *tlx3b*+ dI3 neurons when embryos were heat-shocked at 5 hpf (Fig. 2e). By contrast, the proportion of labeled ventral *foxd3*+ V1 neurons, a ventral subtype not patterned by BMP signaling, remained unchanged. These results demonstrate that BMP signaling acts during gastrulation to specify multiple dorsal interneuron subtypes and reveal decreasing (D-to-V) domain-specific sensitivities to a fixed level of pathway perturbation.

To determine whether this requirement persists beyond gastrulation, we heat-shocked mosaic embryos at 10 hpf, after gastrulation had concluded. Under these conditions, transgenic cells showed a selective reduction in dI1 specification with no significant effect on dI2 or dI3 neurons, consistent with our prior results with post-gastrulation small molecule BMP inhibition (Fig. 2f). This temporal restriction thus supports the conclusion that BMP signaling after gastrulation is required primarily for dI1 specification.

### BMP signaling during gastrulation patterns dorsal interneuron progenitors in a dose-dependent manner

Having established a requirement for BMP signaling during gastrulation, we next asked whether graded levels of BMP activity during this period pattern dorsal progenitor identities in a dose-dependent manner. Using previously defined morphological criteria for dorsoventral pattern disruptions in zebrafish, we identified low (0.2μM) and moderate (0.5μM) doses of DMH1 that permitted reproducible, non-lethal, and graded levels of BMP pathway disruption during gastrulation (Fig 3a)^19,25–27^.

**FIGURE 3.**
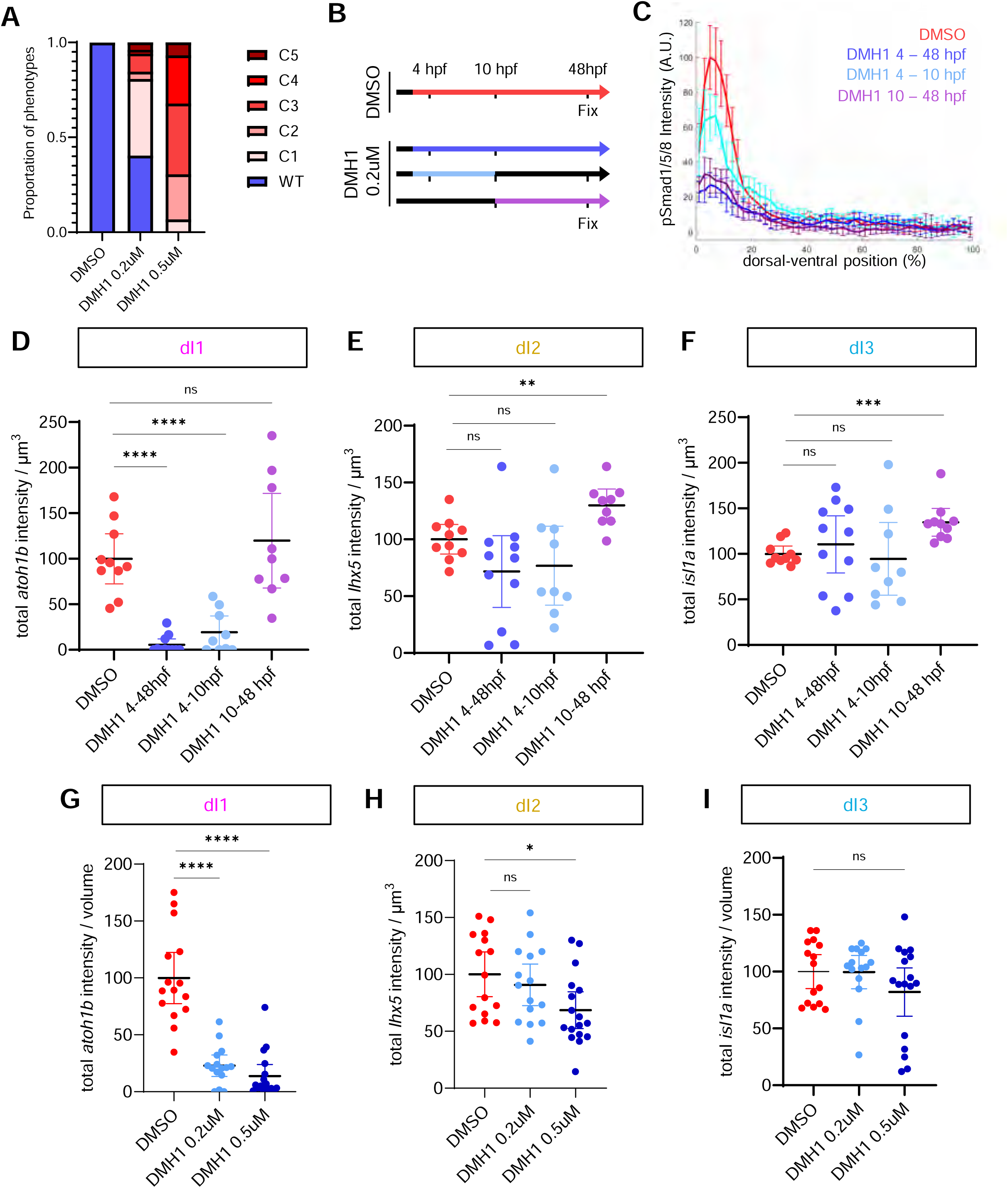
BMP signaling during gastrulation patterns dorsal interneuron identities in a dose-dependent manner. **a** Stacked bar plot of dorsalized phenotypes of embryos treated with either DMSO or 0.2μM and 0.5μM DMH1 from 4 to 24 hpf. (n= 49, 52, 59). **b** Experimental schematic of DMSO or low dose DMH1 treatment. **c** Normalized fluorescence intensity of pSmad1/5/9 immunostaining across of the DV axis of the neural tube after treatment of DMSO or DMH1 (0.2μM) during different temporal windows. (n= 5 each). **d-f** Quantification of the total expression intensity of markers for dI1(*atoh1b)* (d), (*lhx5)* (e), and (*isl1a)* (f) normalized to segmented spinal cord volume after treatment with DMSO or DMH1 (0.2 μM) at the indicated timepoints and fixed at 48 hpf. (n = 10, 11, 9, 10). ****P<0.0001, ***P<0.001, **P<0.01. **g-i** Quantification of the total expression intensity of markers for dI1(*atoh1b)* (g), dI2 (*lhx5)* (h), and dI3 (*isl1a)* (i) normalized to segmented spinal cord volume after treatment with DMSO, DMH1 (0.2 μM), or DMH1 (0.5 μM) from 4-48hpf and fixed at 48 hpf. (n = 15, 15, 17). ****P<0.0001, *P<0.05.

To determine the temporal requirements for BMP-dependent dorsal patterning, we applied low-dose DMH1 (0.2μM) during defined developmental windows before and after gastrulation (Fig. 3b). Continuous low-dose DMH1 treatment from 4–48 hpf resulted in a sustained reduction of pSmad1/5/9 levels within the neural tube, whereas a washout of the inhibitor at 10 hpf largely rescued BMP signaling activity (Fig. 3c). Quantification of dorsal interneurons within a fixed region of the spinal cord showed that continuous low-dose DMH1 treatment selectively reduced the number of dI1 neurons with no significant effect on dI2 or dI3 (Fig. 3d–f, Supplementary Fig. 5a). Similarly, low-dose DMH1 treatment specifically during gastrulation (4-10 hpf) reduced *atoh1b*+ dI1 neurons. Notably, low-dose DMH1 treatment after gastrulation (10–48 hpf) did not reduce any dorsal interneuron subtype and instead increased expression of markers for *lhx5*+ dI2 and *isl1a*+ dI3 neurons, despite a persistent reduction in pSmad1/5/9 levels within the neural tube. Together, these findings identify gastrulation as a critical window during which BMP signaling levels determine dorsal interneuron identity. Analysis of marker expression profiles further revealed that low-dose DMH1 treatment during gastrulation disrupted dorsoventral patterning of dI1 and dI2 neurons, whereas post-gastrulation low-dose DMH1 treatment primarily resulted in dorsal expansion of dI3 markers (Supplementary Fig. 5b–d).

To directly test whether different levels of BMP signaling activity during gastrulation pattern distinct dorsal interneuron identities, we treated embryos with either low or moderate doses of DMH1 continuously from 4–48 hpf (Fig. 3g). Moderate-dose DMH1 treatment (0.5μM) significantly reduced both dI1 and dI2 neurons, whereas low-dose treatment (0.2μM) significantly reduced only dI1 neurons with no effect on dI2 (Fig. 3h, i). Neither the low- nor moderate-dose of DMH1 affected dI3 neurons, consistent with this subtype requiring only minimal levels of BMP signaling activity for specification (Fig. 3j). Together, these results indicate that BMP signaling acts in a dose- and time-dependent manner during gastrulation to pattern multiple dorsal interneuron progenitor identities (dI1 and dI2, respectively), while its later high-level of activity within the neural tube restricts the numbers of dI2 and dI3 neurons.

### The dorsal and ventral spinal cord originate from distinct regions of the gastrula

Our data indicate that BMP signaling during gastrulation is required for dorsal interneuron specification, raising the possibility that these progenitors arise from early embryonic regions exposed to high BMP signaling. To define where dorsal spinal cord progenitors are initially specified, we performed fate mapping using the green-to-red photoconvertible protein Kik-GR^28,29^ (Fig. 4a). One-cell stage embryos were injected with Kik-GR mRNA, and either the ventral or dorsal region of the embryo was targeted with spatially restricted 405 nm photoconversion at the onset of gastrulation (6 hpf) (Fig. 4b, Supplementary Fig. 6a). Embryos were then allowed to develop to 24 hpf and analyzed for the contribution of photoconverted cells to various tissues (Fig. 4c, Supplementary Fig. 6b-d). Control (non-photoconverted) embryos exhibited only green fluorescence, while complete photoconversion produced uniform and stable red fluorescence throughout the embryo (Supplementary Fig. 6b). When only the ventral half of the gastrula was photoconverted, we observed that red fluorescence was stably restricted to ventrally derived tissues, including the epidermis and ventral tail fin^30^ (Supplementary Fig. 6c, d). Conversely, when the dorsal half of the embryo was photoconverted during gastrulation, red fluorescence was restricted to known dorsally derived tissues such as the brain, notochord, and hatching gland (Supplementary Fig. 6c, d).

**FIGURE 4.**
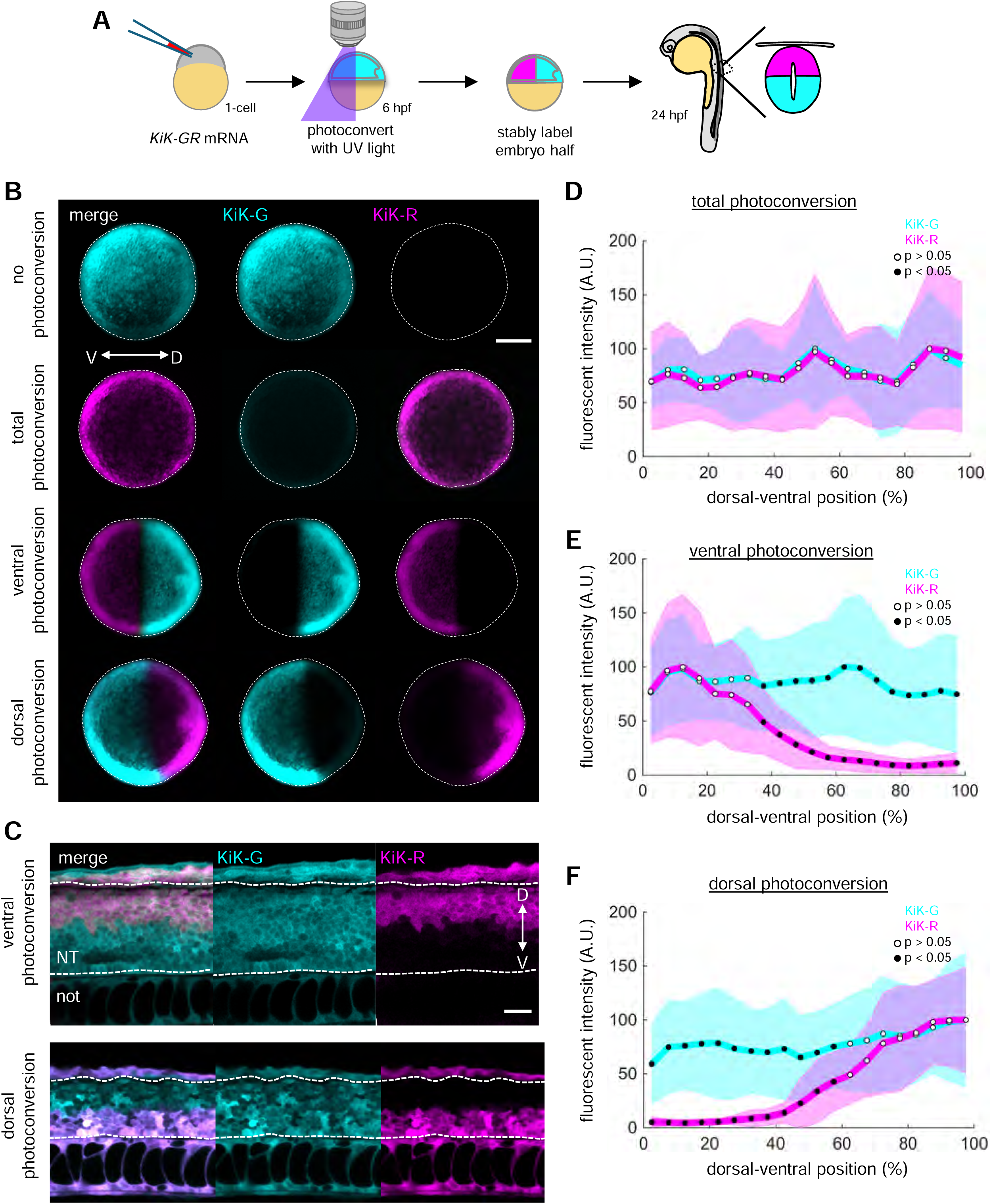
Lineage tracing of dorsal neural progenitors in the gastrula. **a** Experimental schematic of photoconversion-based lineage tracing. Kik-GR mRNA was injected at the 1-cell stage and different regions of the embryo were photoconverted at 5 hpf with 405 nm light. Embryos were evaluated at 24 hpf to determine how photoconverted cells contribute to the spinal cord. **b** Representative images of 6 hpf embryos expressing cytosolic Kik-GR immediately following photoconversion of green to red fluorescence of the entire, the ventral half, or the dorsal half of the embryo. Scale bar, 200 μm. **c** Representative images from a single z-slice of the neural tube at 24 hpf from embryos after ventral or dorsal photoconversion during gastrulation. Scale bar 20 μm. NT, neural tube; not, notochord. **d-f** Average red (photoconverted) and green (unconverted) fluorescence intensity profiles across the DV axis of the spinal cord after total embryo photoconversion (d) (n = 8), ventral photoconversion (e) (n = 26), or dorsal photoconversion (f) (n = 11).

Strikingly, ventrally photoconverted cells contributed exclusively to the dorsal half of the spinal cord, while dorsally photoconverted cells contributed exclusively to the ventral half (Fig. 4c). Fluorescence intensity profiles confirmed this trend: ventral gastrula photoconversion produced strong dorsal enrichment in the neural tube, while dorsal gastrula photoconversion labeled the ventral trunk spinal cord (Fig. 4d–f). This dorsal bias was consistent along the entire anterior–posterior axis (Supplementary Fig. 7a, b, d). We noted that ventral gastrula progenitors contributed to a broader spinal DV domain in the tail posterior to the yolk extension (Supplementary Fig. 7c, e), a result consistent with prior claims of increased vegetal ectoderm contributions to posterior regions of the spinal cord^31^.

These findings demonstrate that dorsal spinal progenitors originate from ventral gastrula cells, an embryonic region exposed to high levels of BMP signaling activity early in development. This establishes a developmental link between the gastrulation BMP gradient and dorsal spinal cord specification, supporting a model in which early BMP signaling pre-patterns dorsal progenitors prior to both neural tube and roofplate formation.

### Progenitors for the dorsal and ventral spinal cord adopt distinct transcriptional states during gastrulation

To determine when dorsal and ventral spinal progenitors are initially specified, and to identify transcriptional regulators of dorsal identity, we interrogated an integrated scRNA-seq atlas (Zebrafish Multi-Atlas Project; ZMAP) comprising 343 single-cell libraries and >750k cells from 8 published zebrafish developmental studies^32^, focusing on neural cells collected from 6 to 10 hpf (Fig. 5a, b). To identify spinal cord progenitors, we calculated a vegetal neural ectoderm (vegNE) marker gene score, which revealed distinct clusters enriched for vegetal markers representing the region where spinal cord progenitors arise (Fig. 5c). We then applied a second gene score to subdivide these vegNE clusters into dorsal and ventral populations based on differential expression of canonical gastrula markers (Fig. 5d). Strikingly, expression of these marker genes revealed an unexpected correspondence between gastrula position and future spinal cord identity. The ventral vegNE cluster, which expressed ventral gastrula markers (*bmp4*, *eve1*), co-expressed dorsal spinal cord markers (*msx1b*, *pax3a*, *olig3*, *zic2a*), while the dorsal vegNE cluster, which expressed dorsal gastrula markers (*chrd*, *admp*), co-expressed ventral spinal cord markers (*nkx6*.1, *olig2*, *ntn1a*) from 6 to 10 hpf (Fig. 5e). This counterintuitive relationship between gastrula position and spinal cord fate is consistent with our lineage tracing results demonstrating that ventral gastrula cells give rise to the future dorsal spinal cord.

**FIGURE 5.**
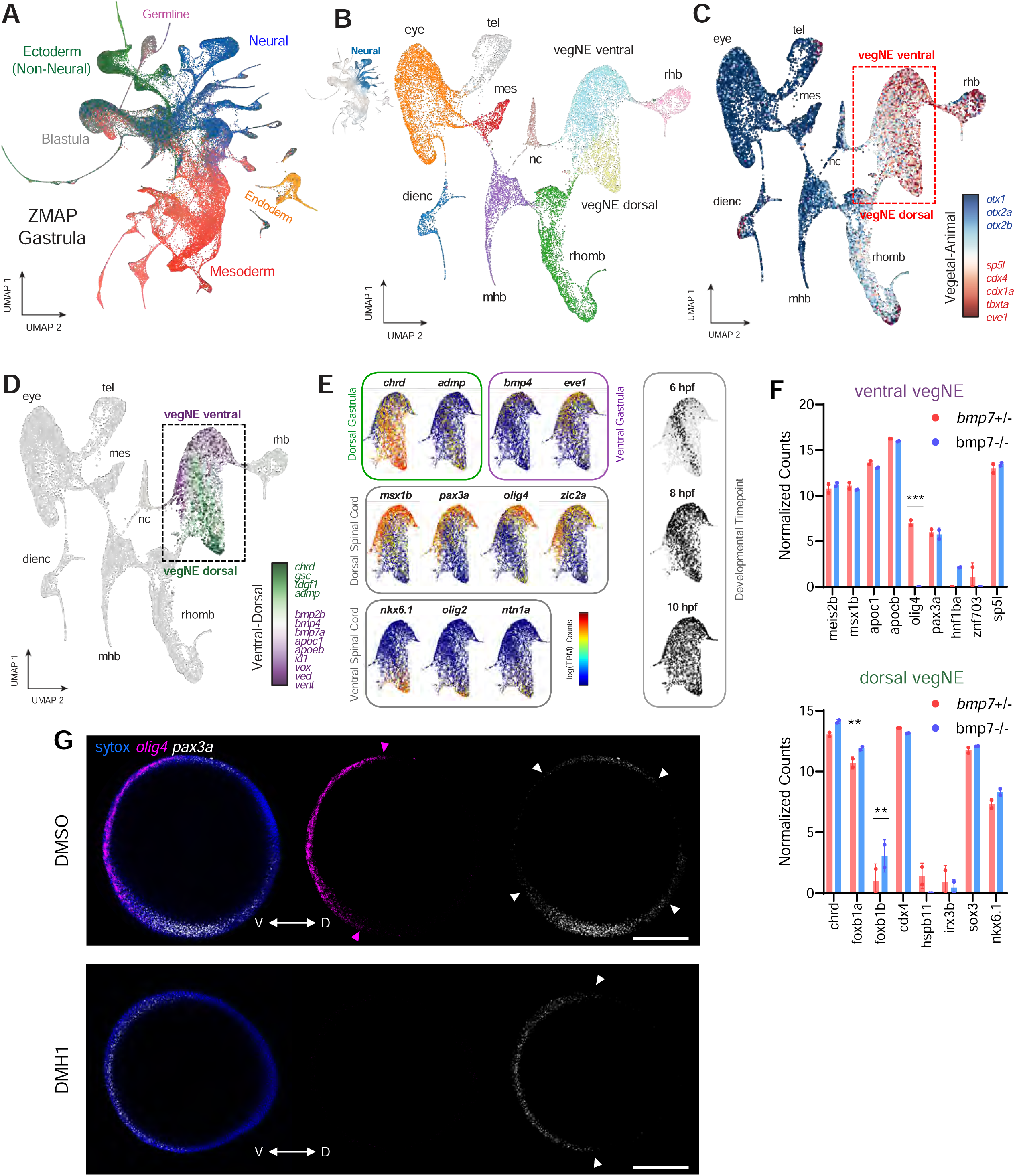
BMP signaling establishes distinct transcriptional states in vegetal neural ectoderm during gastrulation. **a** UMAP embedding of gastrula-stage cells from ZMAP atlas, colored by Tissue-level annotations. **b** Sub-clustering analysis of neurectoderm gastrula cells (cells flagged as Neural in (A). **c** UMAP of neurectoderm clusters, overlaid with Animal-Vegetal gene scores, reveals vegetal neurectoderm (vegNE) cell subset. **d** UMAP of neurectoderm clusters, overlaid with Dorsal-Ventral gene scores, reveals dorsal and ventral subsets of vegetal neurectoderm (vegNE). **e** Inset UMAP overlays of ventral neurectoderm cells depicting expression of dorsal and ventral marker genes for gastrula and spinal cord stages, respectively, and developmental time. **f** Normalized expression from bulk RNA-seq data of *bmp7*+/- and *bmp7*-/- mutant embryos at 70% epiboly for the top ten differentially expressed genes from the ventral and dorsal vegNE clusters. ***FDR < 0.001, **FDR < 0.01. **g** Representative image of HCR *in situ* hybridization for *olig4* and *pax3a* in DMSO- and DMH1-treated (50 μM) embryos at 8 hpf. Arrow heads highlight the boundary of *olig4* and *pax3a* expression. Animal pole is up. Scale bar, 200 μm. (n = 10, 9).

To test whether these cell states required BMP signaling, we analyzed published bulk RNA-seq data from *bmp7*-/- mutant embryos at 70% epiboly^33^ for changes in ventral vegNE and dorsal vegNE markers (Fig. 5f). We found that the ventral vegNE marker *olig4* (*Olig3* in mouse) was significantly downregulated in *bmp7*-/- mutants compared to controls, suggesting that its expression is BMP-dependent. To validate whether BMP signaling regulates expression of *olig4* during gastrulation, we treated embryos with 50 μM DMH1 to abolish BMP signaling activity and examined expression of two markers with opposing BMP dependencies (Supplementary Fig. 8a). HCR analysis revealed that *olig4* was significantly reduced across the dorsoventral axis following DMH1 treatment (Fig. 5g). By contrast, *pax3a*, a vegNE marker negatively regulated by high BMP levels^34^, remained expressed but shifted ventrally into regions normally exposed to high BMP signaling (Fig. 5g). As *olig4* is a key determinant of dorsal interneuron development^13,14,35^, these results identify it as an early BMP-responsive gene linking gastrulation-stage signaling to later dorsal neural fates.

Together, these findings indicate that BMP signaling during gastrulation establishes distinct transcriptional states within the ventral neurectoderm, which initiate dorsal spinal cord programs prior to neural tube formation. This early transcriptional specification provides a mechanistic link between gastrulation-stage BMP exposure and later dorsal progenitor identity, supporting a model in which dorsal neural fate is established during gastrulation rather than within the neural tube.

## Discussion

Our findings fundamentally revise the classic model of dorsal spinal cord patterning. We propose that, rather than functioning as a roof plate-derived morphogen that directly specifies multiple dorsal progenitor identities, BMP signaling is characterized by multiple distinct phases: an early primary phase that pre-patterns dorsal progenitor competence during gastrulation in a dose-dependent manner, and a roof plate-associated phase that maintains dI1 specification and refines neuronal maturation and positioning.

This pre-patterning based framework resolves two major categories of inconsistency between the classic model and decades of experimental data. First, genetic loss of function experiments have consistently revealed markedly reduced dependence of dI2 and dI3 neuronal subtypes on roofplate-associated BMP signaling^7,36–39^. These results have been argued to reflect the redundant activities of the remaining (unperturbed) pathway components. Our work, which shows the same result after severe ablation of post-gastrulation BMP signaling activity, instead indicates that this reduced dependency derives from the time window of patterning rather than genetic redundancy. Second, conditional BMP signaling perturbations have repeatedly revealed evidence of pathway sensitivity that precedes roofplate formation^25,40–42^. These results have generally been explained via reductions in downstream roofplate signaling output. Our work, which explicitly resolves gastrulation and post-gastrulation developmental stages, instead demonstrates that these periods display qualitatively distinct modes of BMP signal interpretation. We propose that gastrulation stage pre-patterning of dorsal neural fates collectively explains these and our findings, and represents a potentially general mechanism for generating neuronal diversity in systems patterned by morphogen gradients.

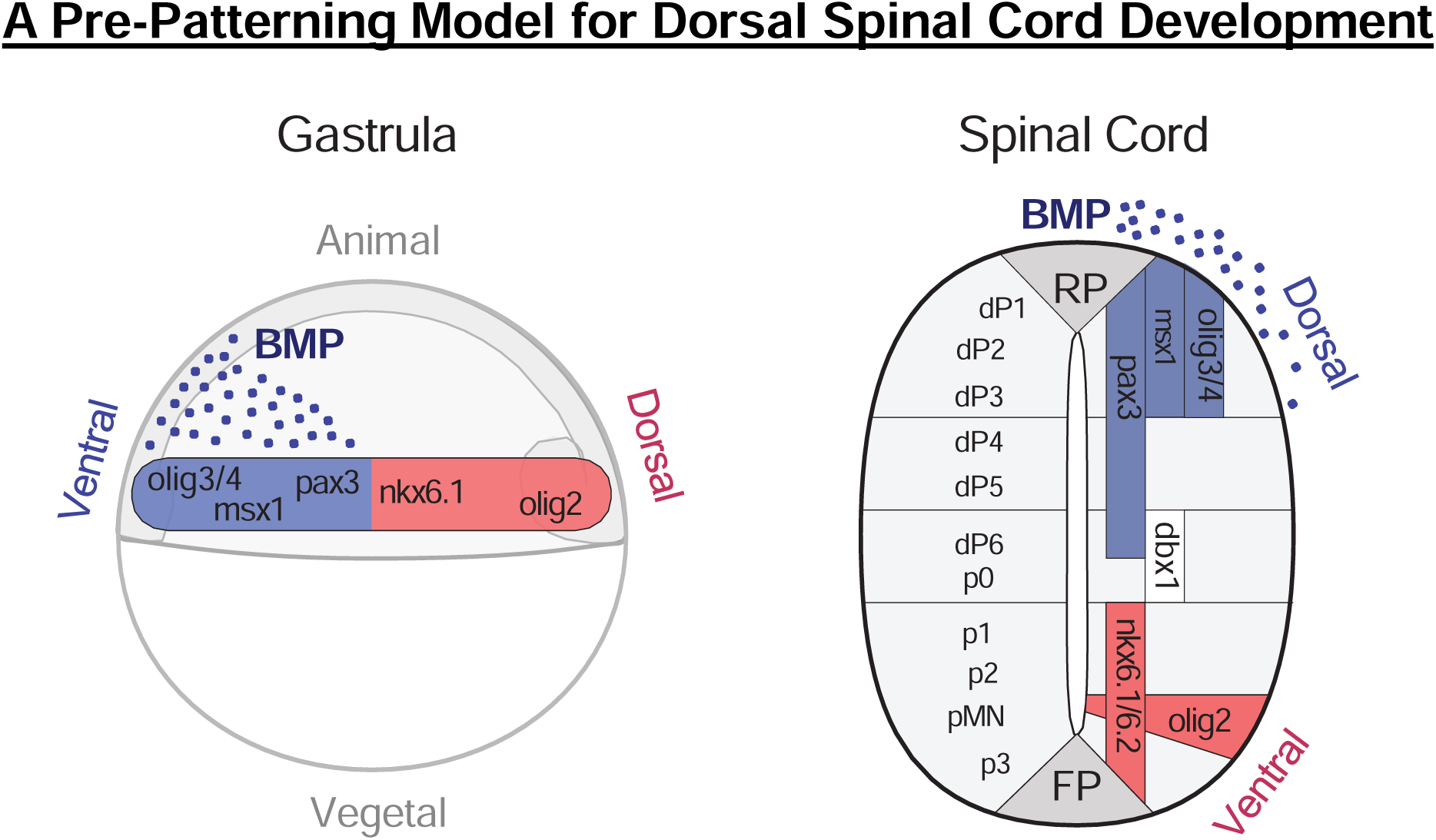

---

**SUPPLEMENTAL FIGURE 1.**
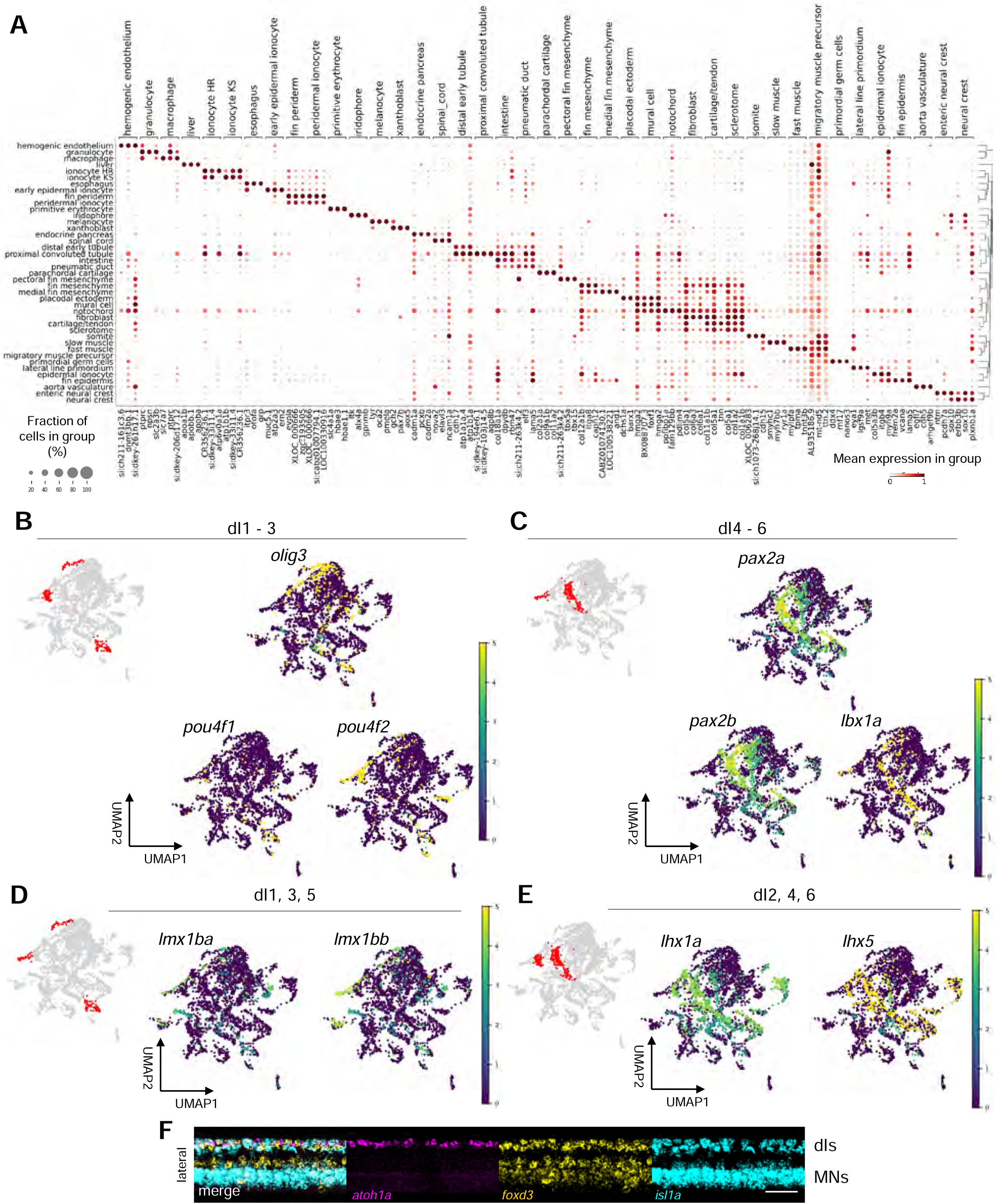
Single-cell RNA sequencing cell type annotation, related to Figure 1. **a** Dot plot depicting the expression of canonical marker genes used to annotate cell types in the 48 hpf trunk scRNA-seq dataset. Dot size represents the percentage of cells expressing the gene, color intensity indicates average expression level. **b-e** UMAP plot of neural cells colored by log-normalized expression of marker genes for dI1-3 (b), dI4-6 (c), dI1, 3, 5 (d), and dI2, 4, 6 (e). Expression of marker gene combinations used to annotate clusters for each dorsal interneuron subtype identities are shown. Inset shows annotated clusters highlighted in red. **f** Full image of the lateral view shown in Fig. 1i of HCR FISH at 48 hpf for all dorsal interneurons dI1 (*atoh1a*), dI2 (*foxd3*), and dI3 (*isl1a*). Scale bar, 30 μm.

**SUPPLEMENTAL FIGURE 2.**
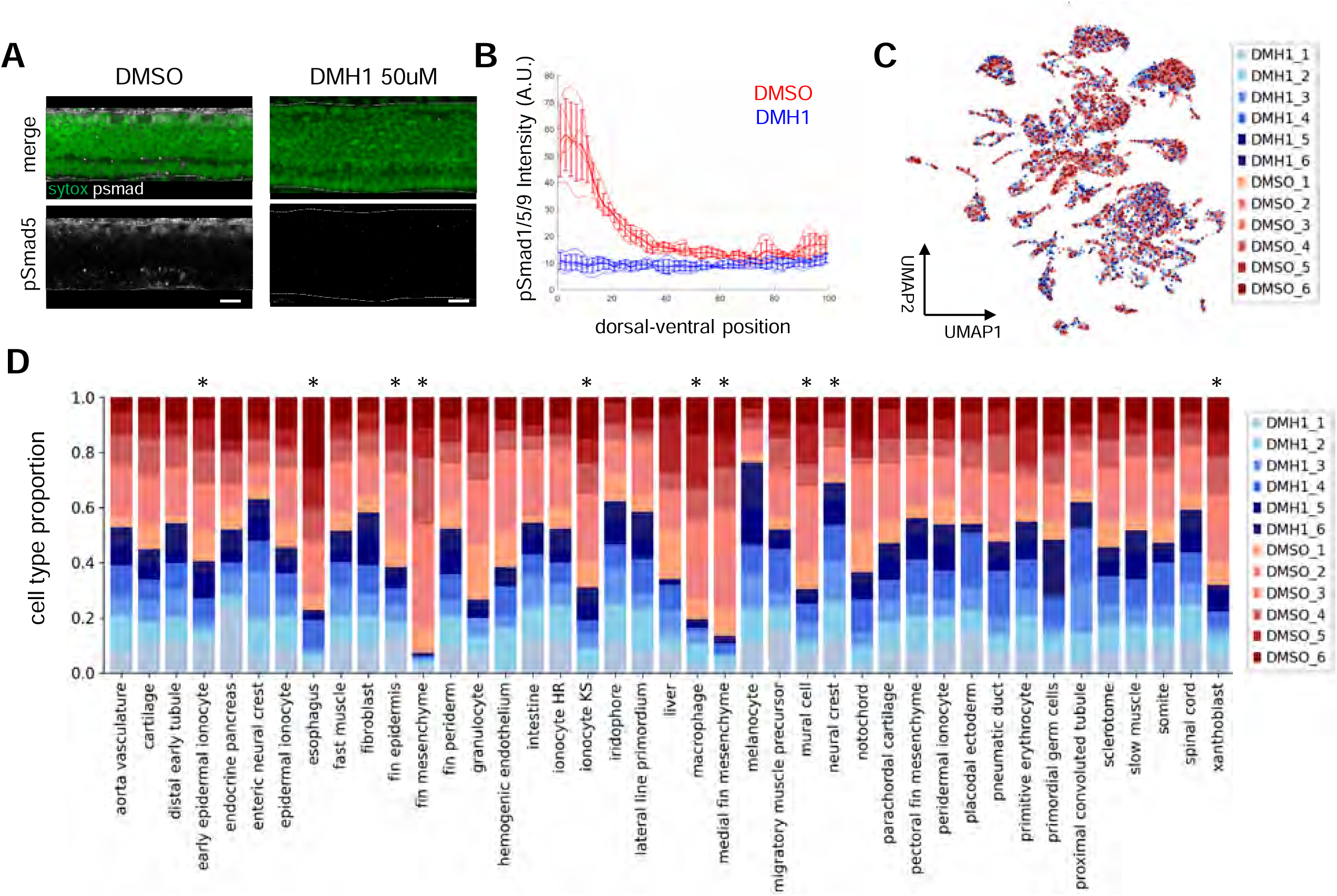
BMP inhibition disrupts cell type proportions during embryonic development, related to Figure 2. **a** Representative images of pSmad1/5/9 immunostaining in 48 hpf embryos treated with DMSO or DMH1 (50 μM) from 10–48 hpf. Scale bar, 50 μm. **b** Quantification of average pSmad1/5/9 intensity across the dorsoventral axis in DMSO- versus DMH1-treated (50 μM) embryos at 48 hpf. (n = 5, 5). (dorsal = 0%, ventral = 100%) **c** UMAP plot of all cells colored by condition and replicate. **d** Differential abundance analysis (see Methods) of cell types between DMSO- and DMH1- treated (50 μM) embryos. Cell types that are significantly differentially enriched or depleted between DMSO- and DMH1-treated embryos are indicated with an asterisk.

**SUPPLEMENTAL FIGURE 3.**
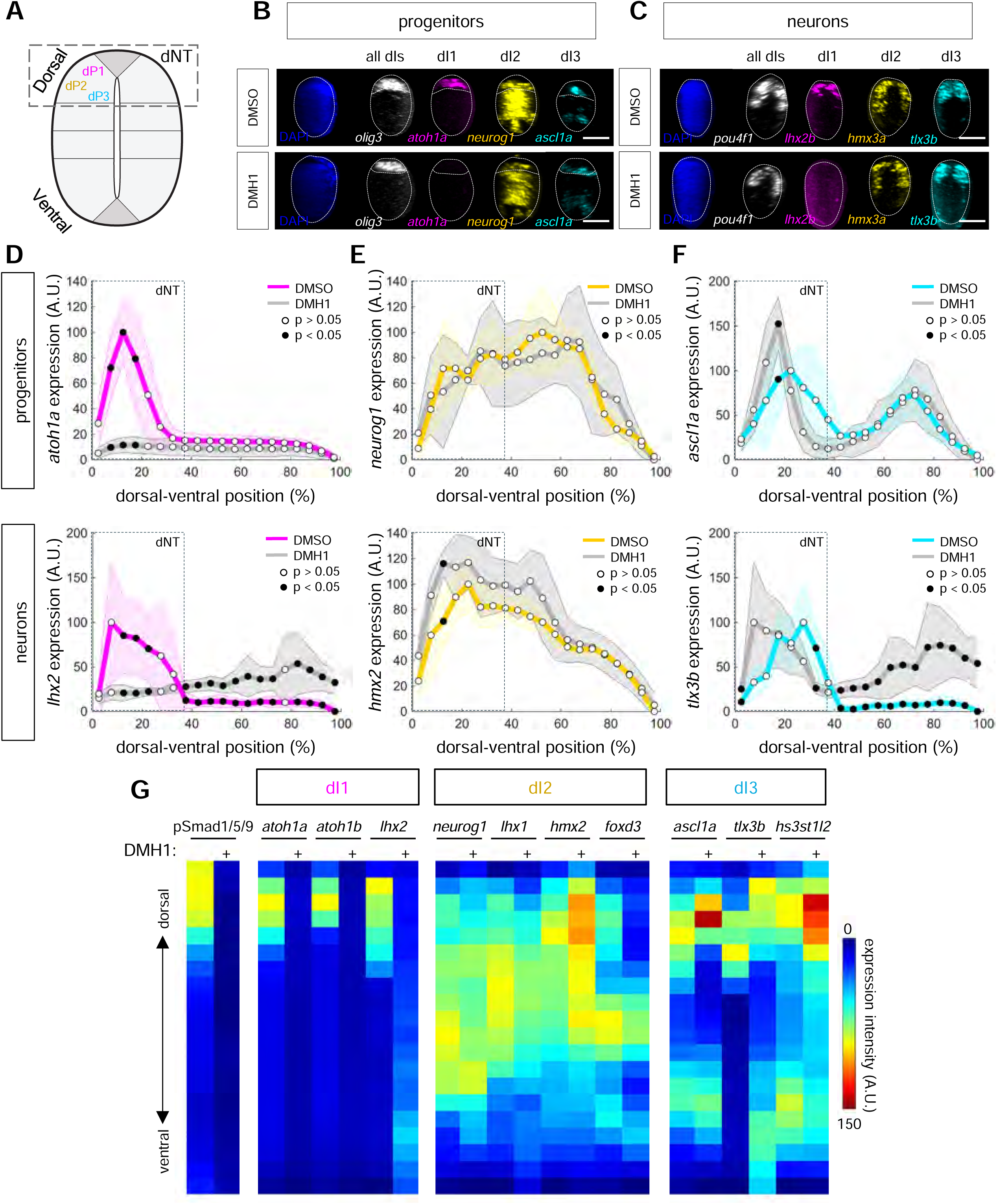
BMP inhibition alters dorsal neural tube composition, related to Figure 2. **a** Schematic of the cell types within neural tube. The dorsal neural tube (dNT) indicated with a box. **b** Representative images of HCR FISH for all dorsal progenitors (*olig3*), dP1 (*atoh1a*), dP2 (*neurog1*), and dP3 (*ascl1a*) in DMSO- and DMH1-treated (50 μM) embryos at 48 hpf. Orthogonal view with dorsal at the top. Scale bar, 25 μm. **c** Representative images of HCR FISH for all dorsal neurons (*pou4f1*), dIN1 (*lhx2b*), dIN2 (*hmx3a*), and dIN3 (*tlx3b*) in DMSO- and DMH1-treated (50 μM) embryos at 48 hpf. Orthogonal view with dorsal at the top. Scale bar, 25 μm. **d** Normalized fluorescence intensities of markers for dP1 (*atoh1a*) and dIN1 (*lhx2b*) across the DV axis of the neural tube at 48 hpf after DMSO (magenta) and DMH1 (gray) treatment (dorsal = 0%, ventral = 100%). **e** Normalized fluorescence intensities of markers for dP2 (*neurog1*) and dIN2 (*lhx1a*) across the DV axis of the neural tube at 48 hpf after DMSO (yellow) and DMH1 (gray) treatment (dorsal = 0%, ventral = 100%). **f** Normalized fluorescence intensities of markers for dP3 (*ascl1a*) and dIN3 (*tlx3b*) across the DV axis of the neural tube at 48 hpf after DMSO (cyan) and DMH1 (gray) treatment (dorsal = 0%, ventral = 100%). **g** Heatmaps of average fluorescence intensities of pSmad1/5/9 immunostaining, and dI1, dI2, and dI3 markers across the DV axis of the neural tube at 48 hpf in DMSO-and DMH1-treated (50 μM) embryos. Expression intensity for each marker is normalized to average maximum intensity (100 A.U.) in DMSO conditions. Blue indicates a decrease in expression, red indicates an increase in expression intensity.

**SUPPLEMENTAL FIGURE 4.**
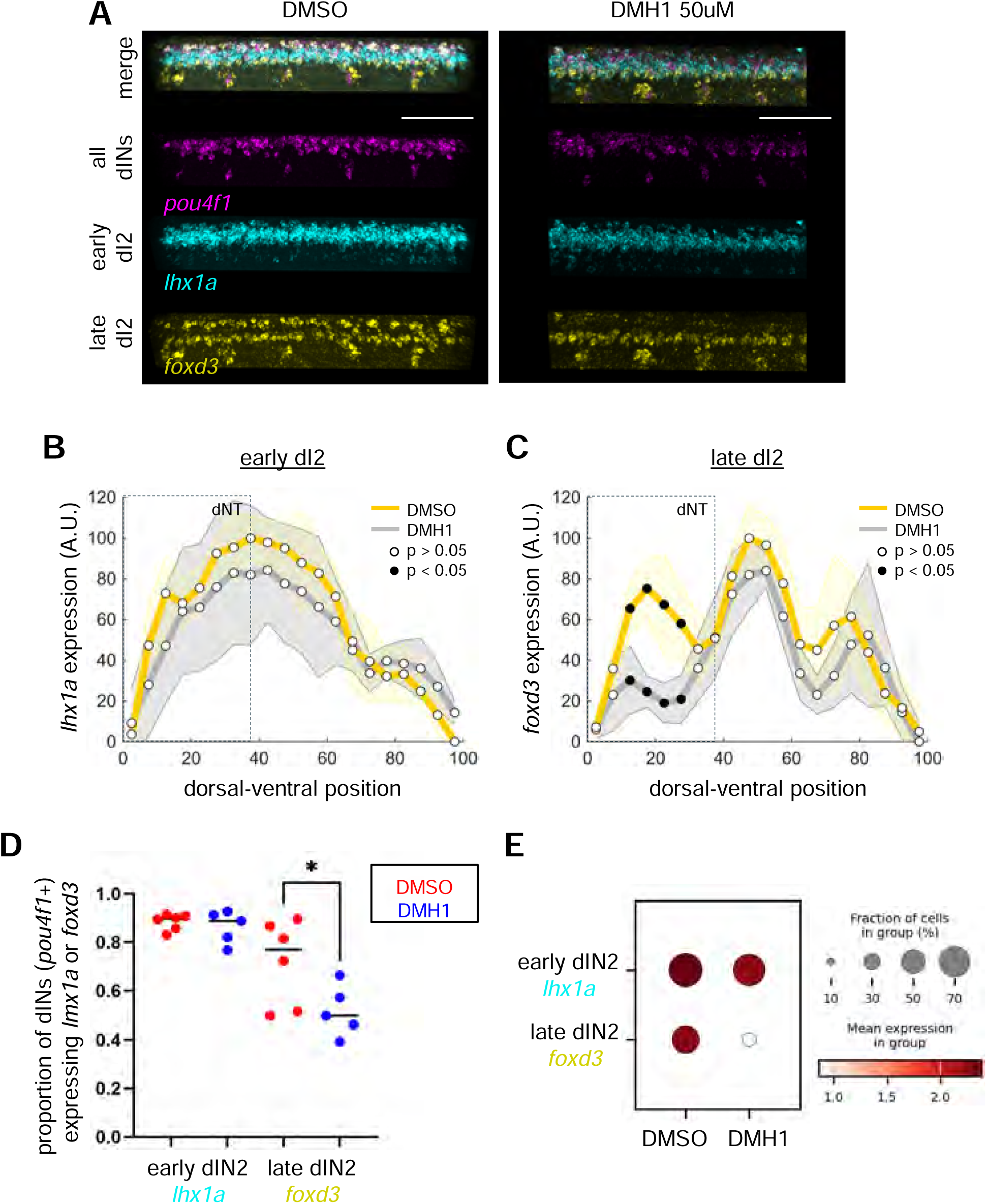
BMP signaling is required for dI2 maturation but not initial specification related to Figure 2. **a** Representative images of HCR FISH of the broad dI marker *pou4f1*, the early dI2 marker *lhx1a,* and the late maturation marker *foxd3* in DMSO- and DMH1-treated (50 μM) embryos at 48 hpf. Anterior to the left and dorsal is up. Scale bar, 100 μm. **b-c** Average DV expression profiles for *lhx1a* (early) and *foxd3* (late). Gray box indicates region of the dorsal spinal cord marked by *olig3* expression. (n = 5 each). (dorsal = 0%, ventral = 100%). **d** Quantification of *foxd3+* vs. *lhx1a+* dorsal interneurons in the spinal cord in DMSO- and DMH1-treated (50 μM) embryos at 48 hpf. *P < 0.05. **e** Dot plot showing average scRNA-seq expression of *lhx1a* and *foxd3* in dI2 cluster cells from DMSO- and DMH1-treated (50 μM) embryos.

**SUPPLEMENTAL FIGURE 5.**
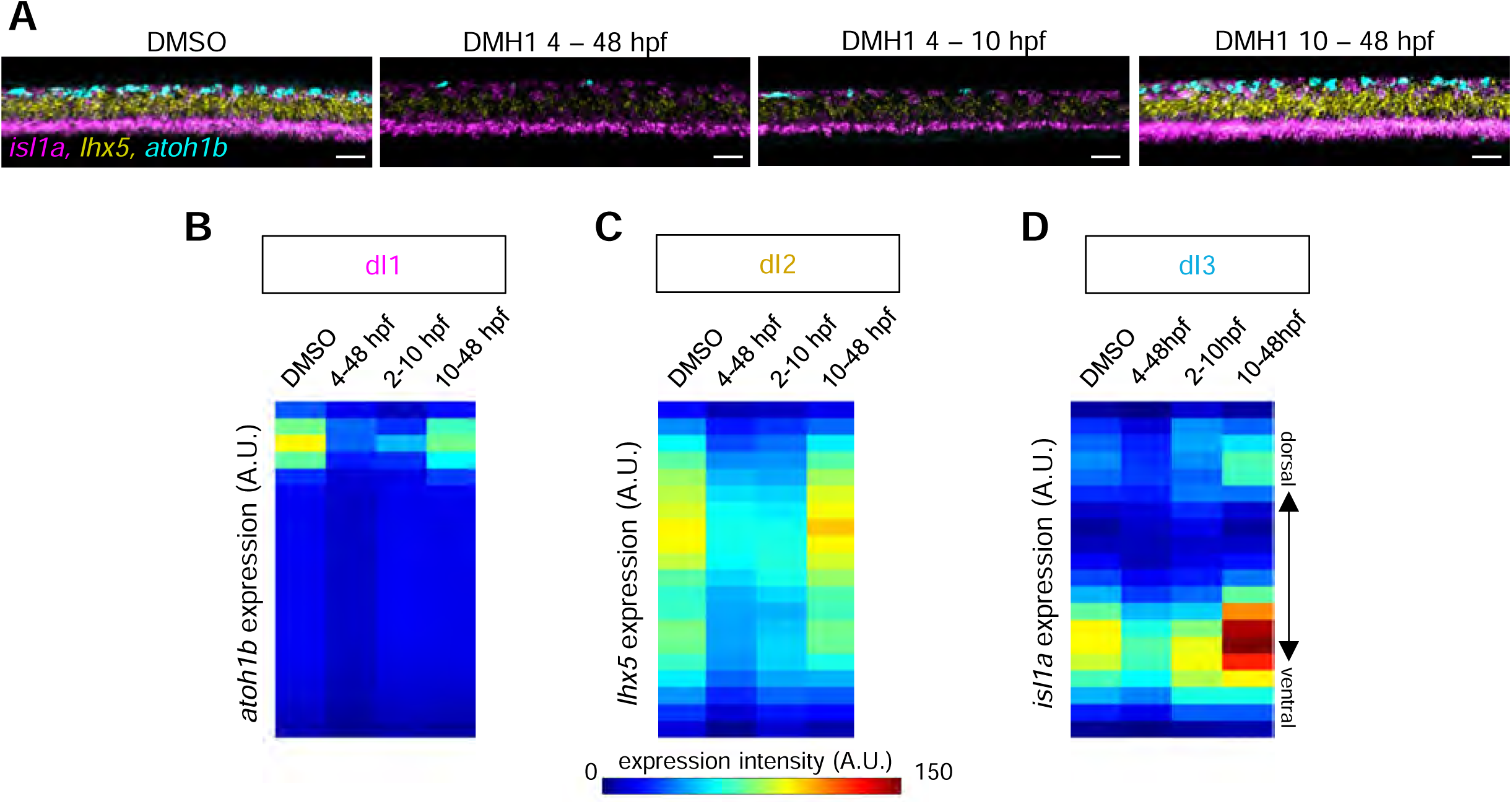
Low-dose treatment of DMH1 reveals critical temporal window during gastrulation, related to Figure 4. **a** Representative images of HCR FISH for markers of dI1 (*atoh1b*), dI2 (*lhx5*), and dI3 (*isl1a*) in DMSO- and DMH1-treated (0.2 μM) embryos at 48 hpf. Lateral view is shown. Anterior to the left and dorsal is up. Scale bar, 100 μm. **b-d** Heatmap of normalized average fluorescence intensity of marker expression across the DV axis of the neural tube at 48 hpf for dI1 (*atoh1b*) (b) (n = 5, 5, 4, 5), dI2 (*lhx5*) (c) (n = 5, 5, 4, 5), or dI3 (*isl1a*) (d) in DMSO- and DMH1-treated (0.2 μM) embryos at 48 hpf (n = 5, 5, 4, 5). Expression intensity for each marker is normalized to average maximum intensity (100 A.U.) in DMSO conditions. Blue indicates a decrease in expression, red indicates an increase in expression intensity.

**SUPPLEMENTAL FIGURE 6.**
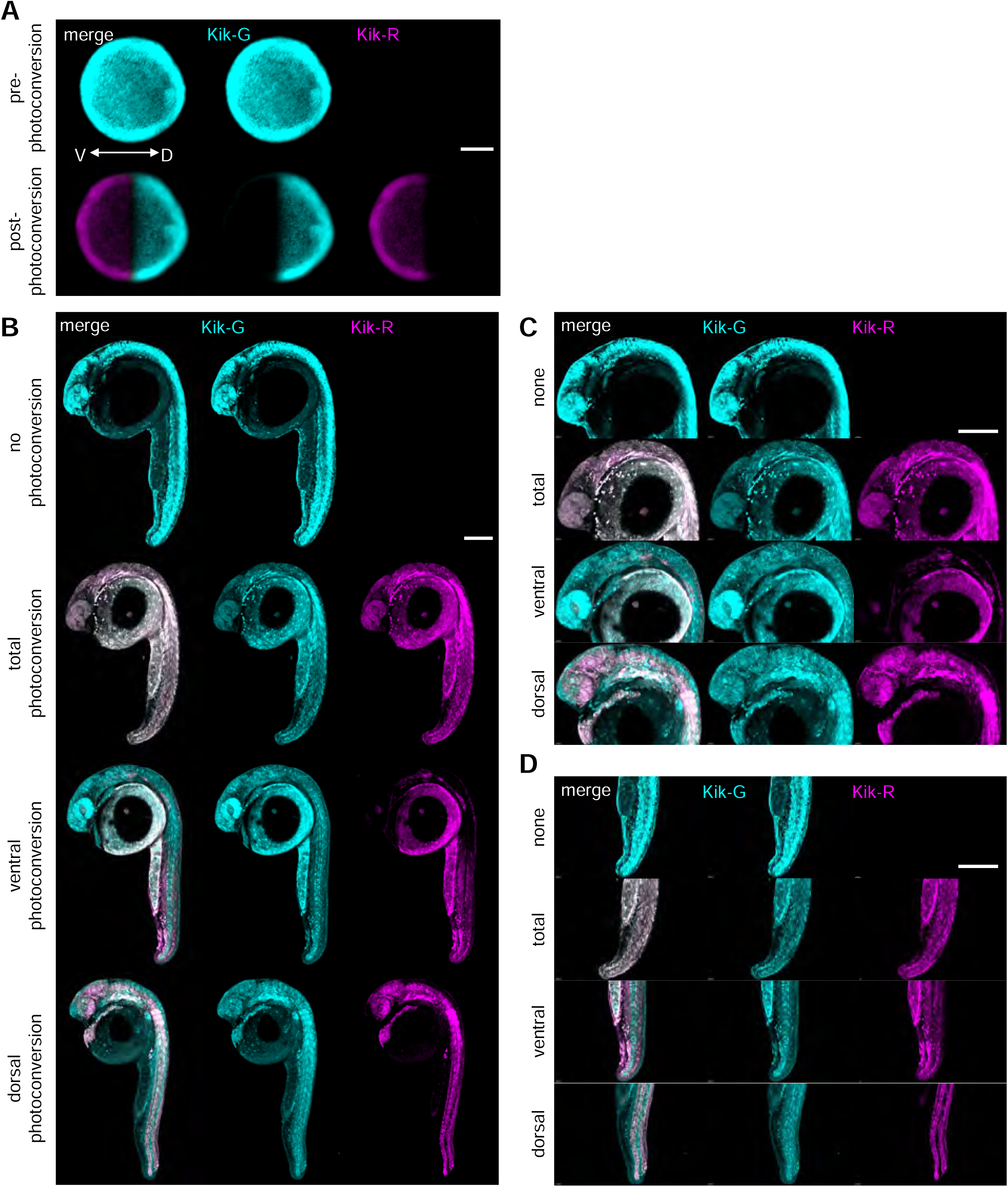
Validation of photoconversion-based lineage tracing, related to Figure 5. **a** Representative images of 6 hpf embryos injected with Kik-GR mRNA pre- (top panels) and post-ventral photoconversion (bottom panels). Animal pole is facing up, ventral is to the left. Scale bar, 200 μm. **b** Representative images of 24 hpf embryos after no photoconversion, total photoconversion, ventral photoconversion, or dorsal photoconversion. Anterior is up, and ventral is to the left. Scale bar, 200 μm. **c-d** Zoomed images showing tissue-specific contributions to the anterior (c) and posterior (d) regions of the embryo at 24 hpf. Scale bar, 200 μm.

**SUPPLEMENTAL FIGURE 7.**
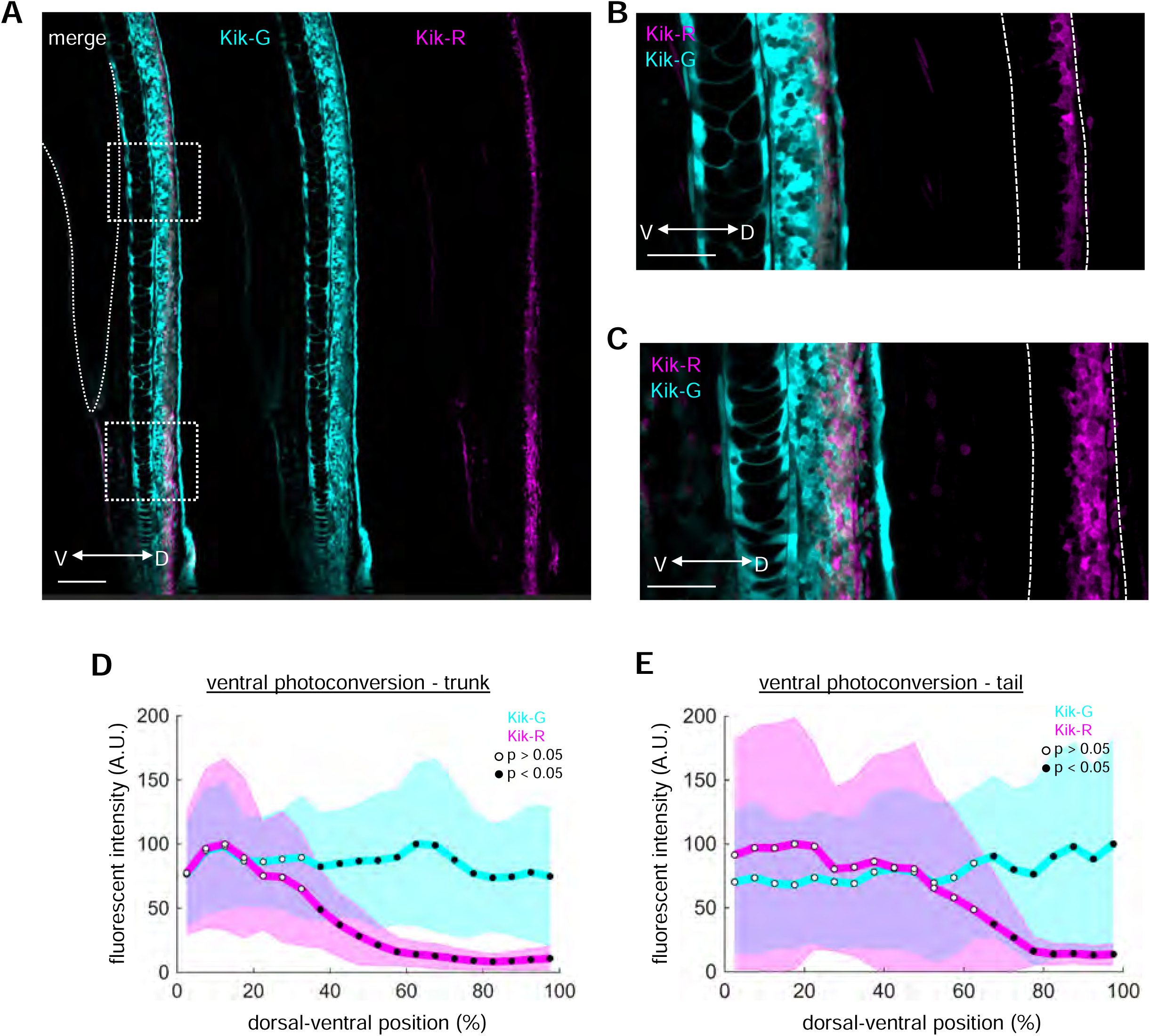
AP-axis distribution of ventrally-derived spinal progenitors, related to Figure 5. **a** Representative images of 24 hpf embryos showing ventral photoconversion contributions along the anterior-posterior axis of the trunk spinal cord Scale bar, 100 μm. **b-c** Zoomed in images of the trunk (b) and tail (c) at 24 hpf after ventral photoconversion. Scale bar, 50 μm. **d-e** Quantification of green and red fluorescence intensity across the DV axis after ventral photoconversion within the trunk (d) (n = 26) and tail (e) (n =10). Shaded regions indicate standard error. (dorsal = 0%, ventral = 100%).

**SUPPLEMENTAL FIGURE 8.**
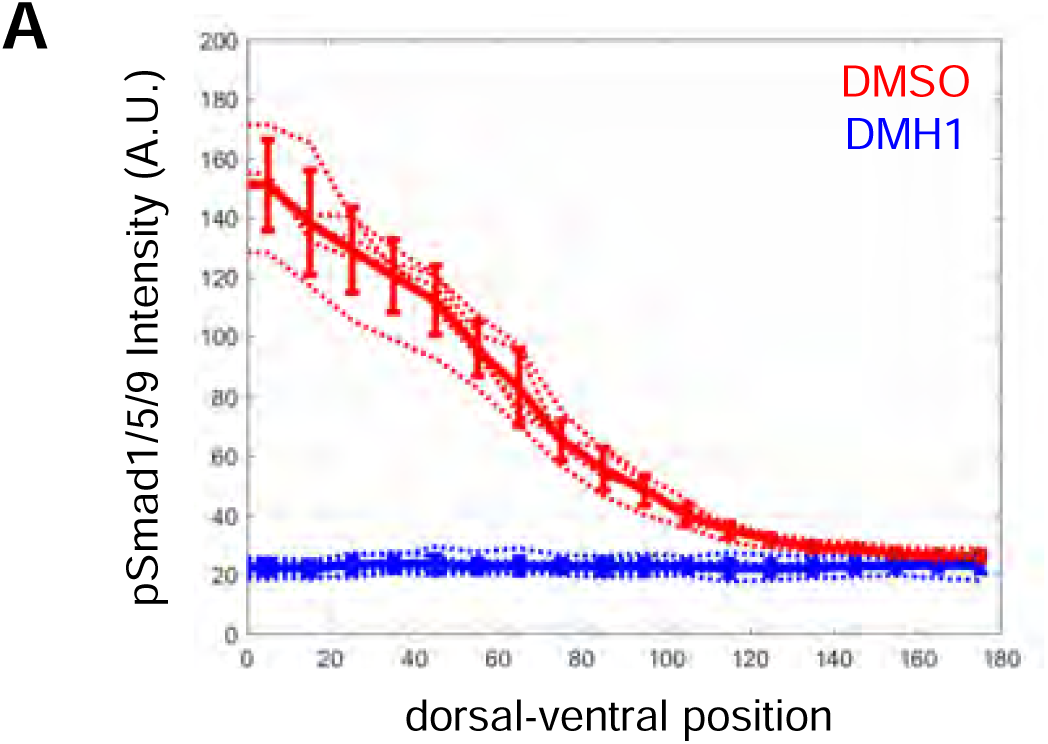
Abolishment of BMP signaling during gastrulation, related to Figure 6. **a** Quantification of average pSmad1/5/9 intensity across the dorsoventral axis in DMSO-versus DMH1-treated (50 μM) embryos at 8 hpf. n = 5 each. (dorsal = 0%, ventral = 100%).

## Methods

### Zebrafish adults and embryos

Adult zebrafish were kept at 28.5 °C in a 14-hour light/ 10-hour dark cycle. All zebrafish husbandry was performed in accordance with institutional and national ethical and animal welfare guidelines. Embryos were collected and raised at 28.5 °C. Sex/gender is not relevant in this study because zebrafish sex determination takes place after 25 days post fertilization ^43^. These studies were performed in accordance with NIH guidelines and those of the University of Californi,a and were approved by the University of California San Francisco IACUC.

### Fluorescent *in situ* hybridization, image processing, and quantification

Zebrafish embryos were fixed overnight in 4% paraformaldehyde at 4°C and gradually dehydrated into methanol. Whole-mount in situ hybridization was performed using hybridization chain reaction v3.0 (HCR3) ^16^. Hybridization mRNA probes, amplifiers, and buffers were purchased from Molecular Instruments, and the protocol provided by the manufacturer for whole-mount zebrafish embryos was followed with no modifications. Embryos were mounted in ProLong Gold Antifade Mountant (Thermo Fisher, P10144) and imaged on an upright Stellaris 5 scanning confocal microscope with a Fluotar 25X water immersion lens. Images were visualized using Imaris to generate 3D renderings (Imaris 9.3.1, Oxford Instruments).

To generate orthogonal views of fluorescent in situ hybridization (FISH) signal within the spinal cord, a 200 µm segment was isolated using Imaris. A surface was rendered based on the DAPI nuclear signal (Life Technologies, D21490) to define the boundaries of the spinal cord in three dimensions. Maximum intensity projections of the defined spinal cord segment were created in FIJI (ImageJ) ^44^ from this surface to visualize the spatial distribution of FISH signal across the dorsoventral axis. For quantification of expression profiles and heatmaps, a straight line was manually drawn from the dorsal-most point to the ventral-most point of the spinal cord maximum projection, ensuring the line spanned the entire dorsoventral axis. The “Plot Profile” function in FIJI was then used to measure the average pixel intensity along this line, providing a quantitative profile of FISH signal distribution across the dorsoventral axis binned into 20 bins. Expression intensity was quantified within a 200 µm segmented region of the spinal cord generated by rendering a surface based on DAPI nuclear signal using Imaris. Total signal intensity was normalized to the volume of the surface.

### pSmad5 immunostaining and quantification

Embryos were fixed in 4% paraformaldehyde at 4°C, blocked in NCS-PBST and probed overnight with a 1:100 dilution of anti-phosphoSmad1/5/9 antibody (Cell Signaling Technology, 13820, RRID:AB_2493181, Danvers, Massachusetts, USA), followed by a 1:500 dilution of goat anti-rabbit Alexa Fluor 647 (Molecular Probes, A-21244, RRID:AB_141663, Eugene, Oregon, USA) and a 1:1,000 dilution of Sytox Green (Thermo Fisher Scientific, S7020, Waltham, Massachusetts, USA). Embryos were mounted and imaged as described above.

Quantification of pSmad1/5/9 within the neural tube was performed as above: a 200 µm region of the spinal cord was segmented by generating a surface rendering using DAPI nuclear signal in Imaris and maximum intensity projected in FIJI. A straight line was manually drawn from the dorsal-most point to the ventral-most point of the spinal cord maximum projection, ensuring the line spanned the entire dorsoventral axis. The “Plot Profile” function in FIJI was then used to measure the average pixel intensity along this line, providing a quantitative profile of pSmad1/5/9 signal distribution across the dorsoventral axis binned into 20 bins.

Quantification of pSmad1/5/9 in gastrula-stage embryos was performed using a published MatLab analysis pipeline ^45^. Briefly, nuclei were segmented by Sytox staining and the pSmad1/5/9 intensity for each nucleus was calculated. Mean intensities for each embryo were calculated and conditions were averaged together for each experimental group. Mean intensities were normalized to the maximum intensity of the control condition. Margin gradients were calculated by isolating a 30um band of cells at the margin in 10-degree bins.

### Inhibition of BMP signaling with a small molecule inhibitor

The BMP signaling inhibitor DMH1 (Fisher Scientific, 2036465MG) was prepared as a 10 mM stock solution in DMSO (Sigma-Aldrich, 41640), aliquoted, and stored at –80°C until use. Aliquots were thawed on the day of each experiment and were not subjected to freeze-thaw cycles or reused. For experimental treatments, DMH1 was diluted in embryo medium to a final concentration of either 50, 0.5, or 0.2µM, depending on the experimental condition. A total volume of 5 mL of the diluted inhibitor solution was dispensed into individual wells of a 6-well plate. Embryos were incubated in inhibitor-containing media at 28°C until fixation at the indicated developmental timepoints. The inhibitor-containing embryo medium was replaced with freshly prepared solution every 24 hours.

### Sample preparation and dissociation for scRNA-seq

Zebrafish embryos were treated with either DMSO or 50 μM DMH1 from 10–48 hpf as described above. At 48 hpf, embryos were anesthetized in 0.2g/L of MS-222 buffered to pH 7.0 (Sigma-Aldrich, A5040), and heads were removed using fine forceps. Twenty trunks per sample were pooled and transferred to low-retention microcentrifuge tubes (Eppendorf, 022431021) on ice.

For single-cell dissociation, samples were incubated in 540 μL of dissociation mixture containing 440 μL of 0.25% trypsin-EDTA (Gibco, 25300054), 60 μL of 100 mg/mL collagenase type I (Sigma-Aldrich, 10269638001), and 40 μL of 1 mg/mL DNase I (Stem Cell Technologies, NC9007308) at 30°C. Tissue was mechanically dissociated by gentle pipetting every 2 minutes until complete disaggregation was achieved (typically 10–15 minutes). Dissociation was quenched by adding an equal volume of pre-warmed DMEM with 10% fetal bovine serum (FBS; Gibco, 16000044). The cell suspension was passed through a 40 μm cell strainer (Corning, 431750), centrifuged at 300 × g for 5 minutes at 4°C, and the supernatant was removed. Cells were washed twice more with wash buffer (1% bovine serum albumin [BSA; Sigma-Aldrich, A7906] in phosphate-buffered saline [PBS]), centrifuging at 300 × g for 5 minutes at 4°C between washes. The final cell pellet was resuspended in 1% BSA in PBS.

Single-cell suspensions were kept on ice, and cell concentration was determined by manual counting using a hemocytometer. Cell viability was assessed by trypan blue (Gibco, 15250061) exclusion. Cells were fixed for split-pool barcoding according to the Parse Biosciences Evercode Cell Fixation v3 protocol (Parse Biosciences, ECF010301). Fixed samples were stored at -80°C until processing.

### Split-pool barcoding and library preparation

Fixed cell samples were thawed on ice and processed using the Parse Evercode Whole Transcriptome (WT) Mini v3 kit (Parse Biosciences, ECW020301) according to the manufacturer’s instructions. A total of 10,000 fixed cells per sublibrary (20,000 cells total across 2 conditions, DMSO and DMH1, with 6 replicates per condition) were loaded into the first round of split-pool barcoding. Split-pool barcoding was performed across three rounds to uniquely barcode individual cells, followed by reverse transcription, second strand synthesis, and PCR amplification.

Library quality and size distribution were assessed using the Agilent Bioanalyzer High Sensitivity DNA kit (Agilent, 5067-4626). Two final sublibraries were generated and sequenced on an Illumina NextSeq 2000 instrument using a P2 flow cell (Illumina, 20046811) with the following read configuration: Read 1 (66 bp), Read 2 (86 bp), Index 1 (8 bp), Index 2 (8 bp).

### Processing of scRNA-seq data

Raw FASTQ files were processed using the Parse Biosciences split-pipe pipeline (version 1.0.3p) and reads were aligned to the zebrafish GRCz11 genome assembly. Gene expression quantification was performed using the annotated GRCz11 gene models, and gene-by-cell count matrices were generated for downstream analysis. Cells were filtered to retain high-quality cells based on the following criteria: >800 UMI counts, and <5% mitochondrial gene content. Doublets were determined and filtered using the Scrublet tool in Scanpy (>0.2 scrublet score). Compositional changes in cell type abundance were assessed using scCODA, implemented in the pertpy Python package and performed using scCODA Hamiltonian Monte Carlo (HMC) sampling with FDR < 0.01 ^46^.

### Phenotypic evaluation

Embryonic phenotypes were assessed at 24 hpf based on previously established criteria: C1 embryos partially lack the ventral tail fin, C2 do not have a ventral tail fin or vein, C3 embryos lack a yolk extension, C4 embryos have small amount of trunk remaining (fewer than 13 somites), and C5 embryos have radialized dorsal somites, seen by elongation at 12 hours post fertilization and lysis by 24 hpf ^19,47^

### Photoconversion lineage tracing

Kikume Green-Red (Kik-GR) photoconvertible fluorescent protein mRNA was synthesized using the mMESSAGE mMACHINE SP6 Transcription Kit (Invitrogen, AM1340) following the manufacturer’s protocol. The synthesized mRNA was purified using SPRIselect (Beckman Coulter, B23317) bead clean-up, and quantified using RNA ScreenTape (Agilent Technologies, TapeStation Analysis Software A.02.01 SRI). One-cell stage embryos were injected with 200 pg of Kik-GR mRNA containing 0.05% phenol red. Injected embryos were maintained at 28.5°C until photoconversion at shield stage (6 hpf).

At shield stage, embryos exhibiting uniform green fluorescence were selected for photoconversion and mounted in 1% low-melting point agarose wells with the animal pole up. Photoconversion was performed on an upright Stellaris 5 scanning confocal microscope using a Fluotar 25X water immersion lens with a 405 nm diode laser at 30% power. The selected region was scanned 100 times (approximately 3 minutes total exposure). Successful photoconversion was confirmed by imaging green (488 nm excitation, 500-550 nm emission) and red (561 nm excitation, 580-650 nm emission) channels before and after photoconversion. Embryos were immediately removed from the wells and maintained at 28.5°C until analysis at 24 hpf.

At 24 hpf, embryos were anesthetized in 0.016% tricaine, mounted laterally in 1% low-melting point agarose in glass-bottom dishes, and imaged on a Stellaris 5 confocal with a 25 and 10X objectives. Z-stacks were acquired with 2 μm step size spanning the entire embryo width. For dorsoventral axis analysis, the neural tube was manually segmented and a straight line ROI was drawn from the dorsal-most to ventral-most point of the neural tube. The line was divided into 20 equal bins and mean red and green fluorescence intensities were measured in each bin using the Plot Profile function. Intensities were normalized to the maximum value within each embryo and profiles from multiple embryos were averaged.

### Mosaic embryo generation and cell-autonomous BMP inhibition

A dominant-negative zebrafish BMP receptor type 1a (dnBMPR1a) construct was generated by cloning the truncated dominant negative form of the Xenopus BMPR1a (Addgene, 15067) ^24^ downstream of the zebrafish heat shock protein 70 (hsp70l) promoter into a Tol2 transposon vector also containing a 2.2kb promoter element from *eef1a1l1* driving H2B-NeonGreen. Tol2 transposase mRNA was synthesized using the mMACHINE SP6 Transcription Kit (Invitrogen, AM1340).

For mosaic analysis, one blastomere of 8-cell stage embryos was microinjected with 25 pg Tol2 transposase mRNA, 25 pg Tg(*hsp*:dnBMPR1-*elf1a*:H2B-NeonGreen) plasmid, and 0.05% phenol red. Injected embryos were maintained at 28.5°C and screened at 6 hpf for mosaic NeonGreen expression. Only embryos showing mosaic labeling (10-30% NeonGreen-positive cells) were retained for heat shock experiments. To induce expression of dnBMPR1, mosaic embryos were heat-shocked at either 5 or 10 hpf by transferring individual embryos to pre-warmed PCR tubes containing 150uL of danieu buffer and heat shocking in the PCR machine set to 37°C for 1 hour. After heat shock, embryos were immediately returned to 28.5°C and maintained until fixation at 48 hpf. Control embryos injected with only efl1a:H2B-NeonGreen plus transposase mRNA were heat-shocked in parallel.

At 48 hpf, mosaic embryos were fixed and processed for HCR *in situ* hybridization for *dnBMPR1* and dIs markers was performed as described above. The number of NeonGreen+dnBMPR1 positive cells were scored for co-expression with neuronal markers within the trunk spinal cord (dI1: *lhx9*, dI2: dorsal *foxd3*, dI3: dorsal *isl1a*). A cell was scored as double-positive only if NeonGreen fluorescence and HCR signal clearly overlapped in single optical sections. For each marker, the percentage of total NeonGreen-positive cells that were also marker-positive was calculated.

### ZMAP gastrula-stage single-cell analysis

The Zebrafish Multi-Atlas Project (ZMAP) integrated single-cell RNA-seq atlas ^32^ was subsetted for all cells through 10 hpf. Cells annotated as neural ectoderm were further subsetted. To identify spinal cord progenitors within the neural ectoderm population, a vegetal neural ectoderm (vNE) gene score was calculated using the ScoreGenes function in Scanpy. To subdivide vNE cells based on their dorsoventral position, two module scores representing dorsal gastrula markers and ventral gastrula markers were calculated also using the ScoreGenes function.

### Analysis of published datasets

Published bulk RNA-seq data from wild-type and bmp7-/- mutant embryos at 70% epiboly ^33^ (GEO accession GSE164760) were analyzed using DESeq2 to identify genes differentially expressed between genotypes.

### Statistical analysis

To determine if two HCR *in situ* expression or pSmad1/5/9 profiles were significantly different, two-tailed T-Tests were performed with a 5% significance level. Profiles shown represent the mean with error bars indicating standard deviation. When performing multiple comparisons, p-values were adjusted using Bonferroni correction for planned comparisons or Benjamini-Hochberg correction for exploratory analyses and RNA-seq, with an FDR threshold of 0.05.

Statistical significance was defined as *p < 0.05, **p < 0.01, ***p < 0.001, or n.s. (not significant, p ≥ 0.05). Error bars represent standard error of the mean (SEM) unless otherwise indicated. Box plots show median (center line), interquartile range (box), 1.5× IQR (whiskers), and outliers (points beyond whiskers). All statistical analyses were performed using GraphPad Prism version 10.0.2.

## Acknowledgments

We thank Y. Su and N. Suren for zebrafish husbandry; C. Chen, K. Chen, C, Chou-Freed, Z. Mendez, I. Roy, N. Aponte Santiago, H. Smith, and J. Zussman, for helpful feedback and discussions.

## Author contributions

Conceptualization: H.G., D.E.W.; Methodology: H.G., D.E.W.; Investigation: H.G., D.E.W.; Visualization: H.G., D.E.W.; Funding acquisition: D.E.W.; Project administration: D.E.W.; Supervision: D.E.W.; Writing – original draft: H.G., D.E.W. All authors reviewed and approved the manuscript.

## Data, code, and materials availability

Raw and processed data will be made available through the NCB1 Gene Expression Omnibus (GEO) and Sequence Read Archive (SRA) repositories, or by request.

