## Supplemental Table 1 for "BMP signaling during gastrulation pre-patterns the dorsal spinal cord"

| Cell Type | Final Parameter | HDI 3% | HDI 97% | SD |
|---|---|---|---|---|
| CoLo | 0 | -0.687 | 0.034 | 0.2 |
| DOLA | 0 | -0.32 | 0.689 | 0.165 |
| KA | 0 | -0.561 | 0.195 | 0.129 |
| MN | 0 | -0.165 | 0.378 | 0.092 |
| OPC | 0 | -0.58 | 0.466 | 0.161 |
| PMn | -0.529990501 | -0.956 | -0.165 | 0.245 |
| Pre-MN | 0 | -0.328 | 0.428 | 0.144 |
| RB | 0 | -1.514 | 0.067 | 0.497 |
| aorta vasculature | 0 | -0.261 | 0.276 | 0.061 |
| cartilage/tendon | 0 | 0.055 | 0.543 | 0.163 |
| dI1 | 1.568352454 | 0.873 | 2.3 | 0.371 |
| dI2 | 0 | -0.819 | 0.13 | 0.265 |
| dI3 | 0 | -1.139 | -0.105 | 0.325 |
| dI4 | 0 | -0.707 | 0.221 | 0.175 |
| dI5 | 0 | 0 | 0 | 0 |
| dI6 | 0 | -0.314 | 0.304 | 0.087 |
| dP | -0.331115966 | -0.53 | -0.154 | 0.113 |
| distal early tubule | 0 | -0.501 | 0.39 | 0.132 |
| early epidermal ionocyte | 0 | -0.016 | 0.882 | 0.301 |
| endocrine pancreas | 0 | -0.674 | 0.695 | 0.267 |
| enteric neural crest | 0 | -0.914 | 0.293 | 0.249 |
| epidermal ionocyte | 0.312530159 | 0.108 | 0.493 | 0.116 |
| esophagus | 1.041823101 | 0.092 | 1.69 | 0.453 |
| fast muscle | 0 | -0.076 | 0.225 | 0.052 |
| fibroblast | 0 | -0.454 | 0.036 | 0.124 |
| fin epidermis | 0.615427195 | 0.368 | 0.883 | 0.146 |
| fin mesenchyme | 1.945975581 | 1.396 | 2.595 | 0.327 |
| fin periderm | 0 | -0.15 | 0.194 | 0.037 |
| granulocyte | 0 | -0.002 | 1.448 | 0.499 |
| hemogenic endothelium | 0 | -0.01 | 1.061 | 0.339 |
| intestine | 0 | -0.56 | 0.383 | 0.141 |
| ionocyte HR | 0 | -0.378 | 0.517 | 0.129 |
| ionocyte KS | 0.804389733 | 0.26 | 1.218 | 0.274 |
| iridophore | 0 | -0.659 | 0.014 | 0.213 |
| lateral floor plate | 0 | -0.568 | 0.429 | 0.121 |
| lateral line primordium | 0 | -0.592 | 0.139 | 0.196 |
| liver | 0 | 0.027 | 1.591 | 0.515 |
| macrophage | 1.369603091 | 0.783 | 2.044 | 0.356 |
| medial fin mesenchyme | 1.790209167 | 1.428 | 2.277 | 0.251 |
| medial floor plate | 0 | -0.505 | 0.333 | 0.162 |
| melanocyte | -0.989978934 | -1.267 | -0.64 | 0.164 |
| migratory muscle precursor | 0 | -0.565 | 0.361 | 0.114 |
| mural cell | 0.833326184 | 0.432 | 1.314 | 0.243 |

| | | | | |
|---|---|---|---|---|
| neural crest | -0.668219282 | -0.97 | -0.31 | 0.182 |
| neural precursor | 0 | -1.064 | -0.156 | 0.322 |
| notochord | 0 | -0.128 | 1.126 | 0.344 |
| oligodendrocyte precursor | 0 | -0.853 | -0.172 | 0.25 |
| p0 | -0.752515027 | -1.228 | -0.326 | 0.247 |
| p2 | 0 | -1.122 | -0.098 | 0.32 |
| pMN | 0 | -0.494 | 0.106 | 0.124 |
| parachordal cartilage | 0 | -0.048 | 0.499 | 0.132 |
| pectoral fin mesenchyme | 0 | -0.478 | 0.154 | 0.162 |
| peridermal ionocyte | 0 | -0.304 | 0.253 | 0.053 |
| placodal ectoderm | 0 | -0.397 | 0.521 | 0.144 |
| pneumatic duct | 0 | -0.269 | 0.61 | 0.167 |
| primitive erythrocyte | 0 | -0.32 | 0.14 | 0.075 |
| primordial germ cells | 0 | -0.559 | 0.843 | 0.234 |
| proximal convoluted tubule | 0 | -0.93 | 0.66 | 0.317 |
| radial glia | 0 | 0.147 | 0.917 | 0.253 |
| sclerotome | 0 | 0.123 | 0.48 | 0.124 |
| slow muscle | 0 | -0.102 | 0.204 | 0.035 |
| somite | 0 | -0.144 | 0.575 | 0.168 |
| v0c | 0 | -1.106 | 0.208 | 0.367 |
| v0v | 0 | -0.577 | 0.181 | 0.174 |
| v2a | 0 | -1.048 | 0.005 | 0.302 |
| v2b | 0 | -0.671 | 0.096 | 0.207 |
| v3 | 0 | -0.671 | 0.708 | 0.169 |
| xanthoblast | 0.831831145 | 0.371 | 1.288 | 0.243 |

| sion proba | ected Sam | log2-fold change | Final Parameter |
|---|---|---|---|
| 0.4268 | 12.26066 | -0.07159845 | FALSE |
| 0.330133 | 3.722502 | -0.07159845 | FALSE |
| 0.288733 | 10.00811 | -0.07159845 | FALSE |
| 0.332267 | 15.88533 | -0.07159845 | FALSE |
| 0.388533 | 4.073066 | -0.07159845 | FALSE |
| 0.960733 | 7.347833 | -0.836213118 | TRUE |
| 0.334267 | 8.153082 | -0.07159845 | FALSE |
| 0.741 | 2.910707 | -0.07159845 | FALSE |
| 0.184133 | 18.14502 | -0.07159845 | FALSE |
| 0.878533 | 28.54255 | -0.07159845 | FALSE |
| 1 | 5.616723 | 2.191055857 | TRUE |
| 0.350533 | 7.311117 | -0.07159845 | FALSE |
| 0.842267 | 8.243261 | -0.07159845 | FALSE |
| 0.339667 | 4.940152 | -0.07159845 | FALSE |
| 0 | 3.913359 | -0.07159845 | FALSE |
| 0.283867 | 12.43352 | -0.07159845 | FALSE |
| 0.970933 | 45.2535 | -0.549297813 | TRUE |
| 0.2844 | 8.571099 | -0.07159845 | FALSE |
| 0.6184 | 5.177882 | -0.07159845 | FALSE |
| 0.5364 | 1.931691 | -0.07159845 | FALSE |
| 0.3938 | 4.064928 | -0.07159845 | FALSE |
| 0.9532 | 70.5221 | 0.37928726 | TRUE |
| 0.9942 | 6.911724 | 1.431434571 | TRUE |
| 0.246 | 116.4423 | -0.07159845 | FALSE |
| 0.325133 | 28.65695 | -0.07159845 | FALSE |
| 0.9964 | 35.79554 | 0.816275312 | TRUE |
| 1 | 12.6085 | 2.73585087 | TRUE |
| 0.1584 | 43.224 | -0.07159845 | FALSE |
| 0.771133 | 1.852241 | -0.07159845 | FALSE |
| 0.7798 | 4.610775 | -0.07159845 | FALSE |
| 0.330467 | 6.510368 | -0.07159845 | FALSE |
| 0.245133 | 5.266658 | -0.07159845 | FALSE |
| 0.997533 | 11.56272 | 1.088890628 | TRUE |
| 0.840333 | 16.04498 | -0.07159845 | FALSE |
| 0.2124 | 4.779787 | -0.07159845 | FALSE |
| 0.4546 | 7.849077 | -0.07159845 | FALSE |
| 0.825533 | 2.480341 | -0.07159845 | FALSE |
| 1 | 8.935205 | 1.904321137 | TRUE |
| 1 | 27.4017 | 2.511127437 | TRUE |
| 0.382933 | 5.722444 | -0.07159845 | FALSE |
| 1 | 10.65913 | -1.499836149 | TRUE |
| 0.233067 | 8.136792 | -0.07159845 | FALSE |
| 0.999733 | 13.16711 | 1.130637103 | TRUE |

| | | | |
|---|---|---|---|
| 0.999667 | 10.22849 | -1.035635094 | TRUE |
| 0.838667 | 11.01652 | -0.07159845 | FALSE |
| 0.6112 | 2.890403 | -0.07159845 | FALSE |
| 0.814867 | 18.21774 | -0.07159845 | FALSE |
| 0.990467 | 5.216783 | -1.157248149 | TRUE |
| 0.831333 | 7.755451 | -0.07159845 | FALSE |
| 0.3002 | 19.15179 | -0.07159845 | FALSE |
| 0.4378 | 16.19004 | -0.07159845 | FALSE |
| 0.3366 | 11.75638 | -0.07159845 | FALSE |
| 0.152333 | 16.22245 | -0.07159845 | FALSE |
| 0.391867 | 4.954994 | -0.07159845 | FALSE |
| 0.4994 | 5.167536 | -0.07159845 | FALSE |
| 0.278067 | 30.67341 | -0.07159845 | FALSE |
| 0.3598 | 2.326572 | -0.07159845 | FALSE |
| 0.421133 | 1.9163 | -0.07159845 | FALSE |
| 0.914867 | 8.700635 | -0.07159845 | FALSE |
| 0.9098 | 50.82539 | -0.07159845 | FALSE |
| 0.143533 | 51.08015 | -0.07159845 | FALSE |
| 0.430133 | 9.888733 | -0.07159845 | FALSE |
| 0.571267 | 2.51531 | -0.07159845 | FALSE |
| 0.524267 | 9.829578 | -0.07159845 | FALSE |
| 0.9072 | 7.325753 | -0.07159845 | FALSE |
| 0.567267 | 10.72305 | -0.07159845 | FALSE |
| 0.222867 | 2.40463 | -0.07159845 | FALSE |
| 0.998933 | 11.12536 | 1.128480218 | TRUE |

**SUPPLEMENTAL TABLE 1**

**Cell type composition analysis in DMSO- and DMH1-treated embryos**

Differential abundance statistics for each cell type are listed. The cell-types that are statistically significantly different between DMSO and DMH1-treated embryos (FDR<0.01) are shown in red.
